# Proteomic Analysis of PTEN-Deficient Cells Reveals Src-Mediated Upregulation of EphA2 and Therapeutic Potential of Dual Inhibition

**DOI:** 10.1101/2025.03.13.643053

**Authors:** Qiong Wang, Xiangyi Kong, Hongming Song, Li Wang, Lingrui Li, Xiaonan Hou, Santosh Renuse, Ran Cheng, Md Kamrul Hasan Khan, Jidong Wang, Kiran Mangalaparthi, Lin Fang, Tamara Levin Lotan, Ben Ho Park, S. John Weroha, Huaijun Zhou, Akhilesh Pandey, Xinyan Wu

**Affiliations:** Department of Molecular Pharmacology and Experimental Therapeutics, Mayo Clinic, Rochester, MN 55905, USA; Department of Obstetrics and Gynecology, Nanjing Drum Tower Hospital Clinical College of Nanjing Medical University; Breast Disease Center, The Affiliated Hospital of Qingdao University, Qingdao, Shangdong 266100, China; Department of Oncology, Division of Medical Oncology, Mayo Clinic, Rochester, MN 55905, USA; Department of Laboratory Medicine and Pathology, Mayo Clinic, Rochester, MN 55905, USA; Department of Obstetrics and Gynecology, Jinan Central Hospital Affiliated to Shandong University, Jinan, Shandong 250013, P.R. China; Department of Breast and Thyroid Surgery, Shanghai Tenth People’s Hospital, Tongji University School of Medicine, Shanghai 200072, China; Department of Pathology, The Johns Hopkins Medical Institutions, Baltimore, MD. 21231, USA; Division of Hematology, Oncology, Department of Medicine, Vanderbilt University Medical Center and the Vanderbilt-Ingram Cancer Center, Nashville, TN, USA; Center for Individualized Medicine, Mayo Clinic, Rochester, MN 55905, USA; Manipal Academy of Higher Education (MAHE), Manipal, Karnataka, India

## Abstract

Loss of the tumor suppressor PTEN is frequently observed in various cancers and promotes tumorigenesis by activating the PI3K-AKT pathway. However, the effectiveness of therapies targeting this pathway is limited by complex signaling crosstalk and compensatory mechanisms. Here, we employed quantitative proteomic and phosphoproteomic analyses using MCF10A PTEN knockout models to comprehensively map the signaling alterations induced by PTEN loss. Our analyses revealed that PTEN deficiency not only activates canonical PI3K-AKT signaling but also induces widespread changes in cytoskeleton organization, cell cycle regulation, and central carbon metabolism. PTEN loss also substantially elevates the activity of a variety of tyrosine kinases, including Src kinase and EphA2, a receptor tyrosine kinase (RTK) implicated in cancer progression. Mechanistic studies demonstrated that Src activation, rather than the canonical AKT signaling pathway, drives the upregulation of the receptor tyrosine kinase EphA2. The activation of the noncanonical tyrosine kinase signaling renders AKT inhibition alone insufficient in PTEN-deficient cancers. Importantly, combined treatment with the FDA-approved AKT inhibitor capivasertib and the Src inhibitor dasatinib synergistically induced apoptosis and suppressed the tumor cell growth in various PTEN-deficient cell lines as well as in three-dimensional cultures of endometrial cancer patient-derived xenograft models. Our study reveals that PTEN loss drives oncogenic signaling via dual activation of PI3K-AKT and tyrosine kinase pathways. Specifically, Src-mediated upregulation of EphA2 in PTEN-deficient cells highlights a therapeutic vulnerability that can be exploited by combined AKT and Src inhibition. This approach addresses the resistance associated with AKT inhibition alone and enhances therapeutic efficacy in PTEN-deficient cancers, supporting its potential application in targeted combination therapies.

## Introduction

PTEN (phosphatase and tensin homolog), a dual-specific phosphatase with both lipid and protein phosphatase activities, acts as a negative regulator of phosphatidylinositol-3-kinase (PI3K) and its downstream signaling cascade (1). PI3K phosphorylates phosphatidylinositol 4,5-bisphosphate (PIP2) to generate phosphatidylinositol 3,4,5-bisphosphate (PIP3), while PTEN antagonizes PI3K by dephosphorylating PIP3 back to PIP2 through its lipid phosphatase activity. In addition to this canonical role, PTEN has protein phosphatase activity and can dephosphorylate serine, threonine, and tyrosine residues, making it a dual-specificity phosphatase (2). PTEN plays a critical role in numerous cellular processes (3) and is frequently defective in a wide range of cancer types (4). For instance, both *PTEN* and *PIK3CA,* which encodes the catalytic subunit of PI3K, exhibit high mutation frequencies across various neoplasms, including glioblastoma, breast cancer, and gynecologic cancers (4). In particular, *PTEN* loss and mutations are observed in >50% of endometrial cancers and can be an early oncogenic event that drives endometrial tumorigenesis through hyperactivation of the PI3K-AKT signaling pathway (5).

PTEN also inhibits cancer initiation and progression through PI3K-independent mechanisms (6). As a protein phosphatase, it regulates cell migration by dephosphorylating FAK in integrin-mediated signaling and SHC in the MAP kinase cascade (7). A translational variant of PTEN, known as PTEN-long, contains extra amino acids at its N-terminal, enabling it to be secreted and reabsorbed by adjacent cells, where it performs diverse functions (8). In addition to its cytosolic activities, PTEN undergoes monoubiquitination followed by translocation to the nucleus (9) where it plays a critical roles in maintaining chromosome integrity (10), DNA damage repair (11), regulating transcription factors (TFs) (12), inducing cell cycle arrest and senescence (13, 14), and interacting with p53 (15). Emerging studies have also highlighted PTEN’s involvement in cancer immune response and cancer metabolism, further underscoring its multifaceted role in tumor suppression (16, 17).

Current treatment strategies for patients with PTEN loss primarily focus on targeting the PI3K-AKT-mTOR signaling pathway. Alpelisib, a PIK3CA inhibitor, was the U.S. Food and Drug Administration (FDA)-approved in 2019 (18), and capivasertib, an AKT inhibitor, in 2023 (19), both in combination with fulvestrant for treating HR-positive, HER2-negative advanced or metastatic breast cancer, with alpelisib targeting PIK3CA mutations (18) and capivasertib addressing PIK3CA, AKT1, or PTEN alterations (19). Clinical trials investigating the potential use of Alpelisib in patients with PTEN loss are ongoing, and favorable results were seen in tumors with no *PIK3CA* alteration but suffering PTEN loss (20, 21). However, prolonged treatment with alpelisib can induce PTEN loss, leading to reactivation of the PI3K pathway and clinical resistance, underscoring the necessity for combination therapies or alternative strategies to overcome this challenge (22).

To uncover novel functions of PTEN and its associated signaling pathways, we conducted Tandem Mass Tag (TMT) labeling-based total proteomic and phosphoproteomic analyses in PTEN-knockout MCF10A cells, revealing significant alterations in pathways including PI3K-AKT-mTOR, MEK-MAPK, cell cycle regulation, senescence, and metabolism. Notably, PTEN loss elevated broad tyrosine kinase signaling, with EphA2, a receptor tyrosine kinase critical for tumor progression, among the most activated, regulated by Src rather than canonical AKT kinase. This highlights the insufficiency of targeting AKT alone in PTEN-deficient cancers, suggesting combined inhibition of AKT and tyrosine kinases like Src as a more effective therapeutic strategy. Supporting this, the combination of capivasertib (AKT inhibitor) and dasatinib (Src inhibitor) synergistically suppressed cancer cell proliferation in PTEN-deficient models, including PDX systems.

This study highlights the complex regulatory mechanisms involving PTEN and its interactions with diverse signaling pathways, paving the way for new opportunities in targeted therapies and combination treatments for PTEN-deficient cancers.

## Experimental procedures

### Cell culture and reagents

All MCF10A cell lines (parental and *PTEN* knockout) were routinely maintained in DMEM/F-12 (1:1) HEPES base medium (Thermo Fisher, 11330032) supplemented with 5% horse serum (HS, Gibco, 26050-088), 20 ng/ml epidermal growth factor (EGF, Sigma), 10 µg/ml insulin (Sigma, I0305000), 0.5 µg/ml hydrocortisone (Sigma, H0888), and 100 ng/ml cholera toxin (Sigma, C8052), along with 100 units/ml Penicillin and 100 µg/ml streptomycin (Gibco, 15140-122). The MCF10A *PTEN* knockout clones were established by homologous recombination (HR)-based somatic gene knockout strategy as described previously (23). Construction of the MCF10A *PIK3CA* (Ex9 and Ex20) knock-in cell lines was described before (24). All MCF10A cell lines were cultured in 0.2 ng/ml EGF as the experimental culture condition. HEK293T, PEO1, MCF7, HCC1937, SPAC-1-L, Ishikawa and HCT116 cell lines were cultured in 10% Fetal Bovine Serum (FBS, Gibco, 26140-079) in RPMI-1640 base medium (Thermo Fisher, 11875093). OV7 was cultured in 5 % FBS in DMEM/F-12 supplemented with 2 mM glutamine, 0.5 ug/ml hydrocortisone, and 10 µg/ml insulin. HEC-1-A was cultured in DMEM/F-12 with 10 % FBS. SNGM was cultured in F-12 HAM’S with 10% FBS. All cell lines were grown in 5% CO_2_ at 37 °C.

To establish doxycycline-induced PTEN overexpression cells, PTEN-deficient SPAC-1-L, HCC1937 and Ishikawa cells were infected with lentiviral particles carrying pLVX-TetOne-PTEN construct. Stably transduced cells were selected using puromycin at concentrations of 5 µg/ml for SPAC-1-L, and 2 µg/ml for both Ishikawa and HCC1937.

### Experimental Design and Statistical Rationale

The MCF10A parental- and two *PTEN* knockout cell lines were cultured in triplicates. TMT labeling and IMAC-based phosphopeptide enrichment were employed for quantitative proteomics and phosphoproteomics analysis. For the phosphotyrosine proteome, SILAC labeling-based experiments were employed. MCF10A parental and *PTEN* knockout cell clones were cultured in DMEM/F-12 SILAC medium without lysine or arginine (Thermo Fisher, 88370) and supplemented with “light-,” “medium-,” or “heavy-labeled” stable isotopic amino acid arginine or lysine (Cambridge Isotope Laboratories) and complete growth factor supplement. The MCF10A parental cell line (“light” label state, L-arginine, L-lysine), *PTEN* knockout cell clone 1 (“medium” label state, L-arginine-^13^C_6_ hydrochloride (Arg +6 Da), L-lysine-^2^H_4_ hydrochloride (Lys +4 Da)), and *PTEN* knockout cell clone 2 (“heavy” label state, L-arginine-^13^C_6_,^15^N_4_ hydrochloride (Arg +10 Da), L-lysine-^13^C_6_,^15^N_2_ hydrochloride (Lys +8 Da)) were maintained in the corresponding media for two weeks before SILAC label check and propagated for the proteomic experiment. MCF10A parental- and two PTEN knockout cell lines were cultured in triplicates for phosphotyrosine proteome analysis. Proteins and phosphorylation sites with significantly differential abundance between experimental conditions were identified by applying a Student’s t test with a p-value <0.05 and fold change greater than 1.5.

To validate these discoveries, multiple cancer cell lines—including breast, ovarian, colorectal, and endometrial—were used in various assays, such as siRNA-mediated knockdown, inhibitor-based kinase activity suppression, TaqMan quantitative RT-PCR, western blotting, cell proliferation assays, and ex vivo three-dimensional culture of patient-derived xenograft (PDX) tumors. Each assay was performed with at least three biological replicates. Statistical significance for all experiments was determined using a Student’s t-test following a normality test. Drug combination synergy was assessed using SynergyFinder (25), and combination indices (CIs) [28] were calculated with CalcuSyn software version 2.1 to evaluate synergy in the ex vivo three-dimensional PDX cultures.

### siRNA-mediated transient knockdown

Transient knockdown of PTEN or EphA2 was performed by the reverse siRNA transfection method. In brief, 30 pmol small interfering RNA (siRNA) and 5 µl of Lipofectamine RNAiMAX Transfection Reagent (Invitrogen, 13778030) were mixed in 500 µl of Opti-MEM medium in 6-cm dishes with 200,000 cells seeded to reach the final siRNA concentration of 10 nM. For 24-well plates, 6 pmol siRNA and 1 µl of RNAiMAX were mixed in 100 µl of Opti-MEM medium to reach the final siRNA concentration of 10 nM. siRNAs include scrambled control (Silencer Select Negative Control, 4390843), siRNAs targeting PTEN (Qiagen, S100301504, S103116092, S103048178, and S100006916), siRNAs targeting Src (Qiagen, S102223921, S102223928) and siRNAs targeting EphA2 (Ambion, s4564 and s4565). Cells were transfected for 72 hours before protein or RNA samples were harvested for the following experiments.

### Kinase inhibition experiments

Kinase inhibitors used in this study, namely alpelisib, MK2206, aapamycin, GSK2256098, dasatinib, U0126, trametinib, ipatasertib and capivasertib, were all purchased from Selleckchem. Cells were treated with inhibitors for 24 hours prior to protein collection except for the time series analysis. For MCF10A, cells were first seeded with a normal medium containing 20 ng/ml EGF. After the cells were well attached to the culture plate surface, they were washed with PBS once, and the cell medium was replaced by inhibitors prepared in 0.2 ng/ml EGF medium.

### Western blot

For immunoblotting, cells were harvested and lysed in modified RIPA buffer (50 mM Tris-HCl with pH 7.4, 1% Nonidet P (NP) −40, 0.25% sodium deoxycholate, and 150 mM NaCl, supplemented with protease and phospho-protease inhibitors). Cell lysate was sonicated to achieve a complete lysis, followed by centrifugation at 20,000 x g for 10 min. Protein supernatants were quantified with BCA protein assay (Pierce^TM^, 23227), denatured at 70 °C in sample buffer (Invitrogen) supplemented with β-mercaptoethanol, and separated in NuPAGE gels (Invitrogen). Afterward, gels were transferred to nitrocellulose membranes and probed with the primary antibodies, followed by incubation with horseradish peroxidase-conjugated secondary antibodies. Anti-PTEN (22034-1-AP) was purchased from Proteintech. Anti-AKT pS473 (4060T), anti-total-AKT (4691S), anti-ERK1/2 pT202/Y204 (4370S), anti-total ERK1/2 (4695S), anti-EphA2 (6997S), anti-SPHK1 (12071), anti-DKK1 (48367), anti-TGFBR2 (41896), anti-Rab27B (44813), anti-Src_pY416 (6943S), anti-total-Src (2109S), anti-P70S6K_pT389 (97596S), anti-GSK-3α/β pSer21/9(9331S), anti-PARP(9542T), Caspase 3 (14220T), Cleaved Caspase 3 (9664T), and anti-actin (4970L) antibodies were purchased from Cell Signaling Technology (CST). Anti-GSK-3α/β (sc7291) was purchased from Santa Cruze.

### Quantitative Reverse Transcriptase PCR (RT-qPCR)

Total RNA was extracted from MCF10A cells using the Monarch® Total RNA Miniprep Kit (T2010S) following the vendor’s protocol. The concentration of sample RNA was assayed by NanoDrop One (Thermo Fisher). cDNA was synthesized using First-Stand buffer (5X) and SuperScript^TM^ III RT (Invitrogen, 18080085). cDNA samples were mixed with the TaqMan® Universal PCR Master Mix (Applied Biosystems, 4304437), EphA2 (Thermo Fisher, Hs01072272_m1, FAM) and actin (Hs99999903_m1, VIC) TaqMan primers, and RNase-free water for RT-qPCR that was carried out in QuantStudio 3 (Thermo Fisher). High ROX dye (50x, Qiagen) was added for normalization.

### Crystal violet proliferation assay

Cells were washed once with PBS before adding 250 µl of 0.02% crystal violet in 5% formalin to completely cover each well of the cell culture plate, followed by incubating at room temperature for 10 min. The crystal violet solution was disposed of, and the cell plate was washed with water to remove excess crystal violet solution. For quantification, the cell-stained crystal violet was eluted with 200 µl of 3% acetic acid, and the well-dissolved solution was measured in a 96-well plate at 570 nm (200 µl). The drug combination synergy was calculated using SynergyFinder (25).

### Establishment of EC PDXs

Under the aegis of protocols approved by the Mayo Clinic IRB and Animal Care and Use Committee, tumor specimens from consenting EC patients obtained during primary surgery or clinical biopsy for recurrent disease were minced in McCoy’s medium, supplemented with penicillin/streptomycin and rituximab (10 mg/kg) (Rituxan; Genentech, Inc., San Francisco, CA) to prevent unintentional lymphoproliferative tumors (26), and injected intraperitoneally into female SCID-bg mice (C.B.-17/IcrHsd-Prkdcscid Lystbg; ENVIGO). Mice were monitored weekly for tumor engraftment and euthanized when moribund criteria were met (27). Minced tumors were cryopreserved for subsequent studies as a 1:1 suspension in freezing medium (39% FBS, 10% dimethyl sulfoxide, 1% penicillin/streptomycin in McCoy’s medium). Key clinical characteristics of the 3 PDX tumors with functional assessment of homologous recombination (HR) activity are described in Supplemental data 9.

### Ex vivo Tumor 3D culture

Viable tissue of EC PDXs were dissociated with the gentleMACS™ Dissociator and plated in ultra-low attachment 384 well microplates (CLS3571, Corning Life Science, USA) in DMEM supplemented with 15% FBS. After 24 h, dasatinib (0.25, 0.125, 0.0625 or 0.03125 mM), capivasertib (1, 0.75, 0.5 or 0.25 mM), or the combination (at the same concentrations) were added to triplicate wells. After 72 hrs, response was determined by the RealTime-Glo MT Cell Viability Assay (G9711, Promega Corporation, USA) in GloMax Discover System (GM3000, Promega Corporation, USA).

### Combination Index Calculations

To determine the synergy in PDX ex vivo 3-D cultures, combination indices (CIs) (28) were calculated using CalcuSyn software, v2.1 (Biosoft, Cambridge, UK) under the assumption that effects are mutually exclusive, which yields results comparable to isobologram analysis (29). CI>1 indicates antagonism, CI=1 indicates additivity, and CI <1 indicates synergy. For RealTime-Glo assays, fraction affected indicates mean decrease in signal compared to the control.

### TMT labeling-based total proteomics and phosphoproteomics

#### Protein extraction, in-solution trypsin digestion, and desalting

The MCF10A parental- and two *PTEN* knockout cell lines were cultured in triplicates, then harvested and lysed in 8 M urea buffer (8 M urea, 20 mM HEPES pH 8.0, 1 mM sodium orthovanadate, 2.5 mM sodium pyrophosphate, 1 mM β-glycerophosphate, and 5 mM sodium fluoride). Cell lysates were then sonicated and subsequently centrifuged at 15,000× g for 20 min at room temperature to eliminate insoluble cell debris. The protein supernatants were quantified with BCA protein assay. 2 mg proteins per sample were used for in-solution trypsin digestion where protein samples were reduced with 5 mM dithiothreitol (DTT) at 37 °C for 1 h, followed by alkylation with 10 mM iodoacetamide (IAA) protected from lights at room temperature for 30 min. The reduced and alkylated protein samples were then diluted 4 times by 20 mM HEPES buffer (pH 8.0) to reach a final urea concentration of less than 2 M. After dilution, protein samples were in-solution digested by protease TPCK-treated trypsin (1:20 (w:w), Worthington Biochemical Corp. Lakewood, NJ, USA) at room temperature overnight on an orbital shaker. On the following day, the digestion reaction was first quenched by adding 20% trifluoroacetic acid (TFA) into the peptide digests to a final concentration of 1% TFA to acidify the reaction environment. The peptide samples were then centrifugated at 10,000× g for 10 min at room temperature. Peptide samples were desalted by SepPak C18 cartridge (Waters Corporation, Milford, MA, USA), and eluted with 0.1% TFA in 60% acetonitrile (ACN). The eluted peptides were vacuum-dried in SpeedVac (Thermo Fisher Savant, SPD120).

#### TMT labeling

Samples from speed vacuum were dissolved in 50 µl of 100 mM triethylammonium bicarbonate (TEABC), and the peptide colorimetric assay was used to detect the peptide concentration (Thermo Fisher, 23275). 1 mg of each peptide sample was then mixed with the TMTpro Label Reagent (Thermo Fisher) dissolved in anhydrous acetonitrile. The mixtures were then incubated at room temperature for 1 hour, and a TMT labeling check was conducted. The labeling reaction for each sample was quenched by adding 10 µl of 5% hydroxylamine and incubated for 15 min at room temperature. TMT reagents-labeled peptides were then mixed into one sample and desalted with C18 reverse-phase column. Desalted peptides were eluted with 0.1% TFA in 60% ACN and lyophilized to dryness. The lyophilized TMT-labeled peptide mixture was kept at −80 °C prior to fractionation and immobilized metal affinity chromatography (IMAC)-based phosphopeptide enrichment.

#### Basic Reversed-Phase Liquid Chromatography (bRPLC) fractionation

The bRPLC fractionation procedure followed the previously described protocol(30)., the lyophilized TMT labeled peptides were reconstituted in 7 mM TEABC (pH 8.5) and fractionated by bRPLC chromatography on a C18, 250 × 4.6 mm column, 5 µm, XBridge (Waters, Milford, MA) by employing an increasing gradient of bRPLC solvent B (7mM TEABC, pH 8.5, 90% acetonitrile) on an Agilent 1260 HPLC system. A total of 96 fractions were collected and combined into 24 fractions.10% of each fraction was used for LC-MS/MS-based for total proteome analysis. The rest of the 90% was vacuum-tried and later subjected to IMAC-based phosphopeptide enrichment.

#### IMAC-based phosphopeptide enrichment

IMAC beads charging with FeCl_3_ were prepared by stripping nickel from Ni-NTA Superflow agarose beads (Qiagen, 1018611) with 100 mM EDTA and incubated in an aqueous solution of 10 mM FeCl_3_ (Sigma, 451649) and 0.1% TFA in 80% ACN. The dried peptide fractions were reconstituted in 0.1% TFA in 80% ACN to achieve a final concentration of 0.5 µg/µl. IMAC beads were added into each fraction (10 µl IMAC beads for 1 mg peptides), and the mixtures were incubated on an end-over-end rotator at room temperature for 1 hour. After enrichment for phosphorylated peptides, IMAC beads with phosphopeptides were loaded on stage tips packed with Empore C18 silica, followed by washing with 1% TFA in 80% ACN. 500 mM dibasic sodium phosphate, pH 7.0 (Sigma, S9763) was used for phosphopeptides elution from IMAC beads.

#### Liquid chromatography (LC)-MS/MS analysis

Fractionated peptides and enriched phosphopeptides were vacuum-dried and reconstituted in 0.1% formic acid and were analyzed on a reverse-phase LC system interfaced with an Orbitrap Fusion Lumos Mass Spectrometer. Peptides were separated on an analytical column (75 µm × 50 cm, RPLC C18) using a 100 min gradient from 8–50% acetonitrile in 0.1% formic acid at a flow rate of 0.3 µl/min. The total run time was set to 120 min. The spray voltage was set to 2.1□V while the capillary temperature was set to 200L°C. The mass spectrometer was operated in a data-dependent acquisition mode. Precursor MS scan (from m/z 300–1700) was acquired in the Orbitrap analyzer with a resolution of 120 000 at 200 m/z. The AGC target for MS1 was set as 1 × 10^6^ and ion filling time set 100 ms. The most intense ions with charge state ≥2 were isolated in 3 s cycle and fragmented using HCD fragmentation with 35% normalized collision energy and detected at a mass resolution of 30k at 200 m/z. The AGC target for MS/MS was set as 5 × 10^4^, and the ion filling time was set to 100 ms. Dynamic exclusion was set for 40 s with a 7 ppm mass window.

#### Mass spectrometry data analysis

The Proteome Discoverer software suite (v 2.5; Thermo Fisher Scientific) was used for the downstream protein identification and quantification. The acquired MS/MS data were searched by the SEQUEST search algorithm against the Human Uniport protein database (20,395 protein sequences, ver. 02012021) supplemented with frequently observed contaminants. The following search parameters were used: (1) trypsin as a protease with full specificity; (2) a maximum of 2 missed cleavages allowed; (3) fixed modification: carbamidomethylation of cysteine and TMTpro tag (+229.163 Da) on lysine residues or peptide N-terminus; (4) variable modification: N-terminal acetylation, oxidation at methionine, and phosphorylation at serine/threonine and tyrosine; (5) precursor tolerance was set at 10 ppm; (6) the fragment match tolerance was set to 0.02 Da.; (7) the PSMs, peptides and proteins were filtered at the cut-off 1% false discovery rate (FDR) calculated using target-decoy database searches. The probability of an identified phosphorylation of specific Ser/Thr/Tyr residue on each identified phosphopeptide was determined using the PhosphoRS algorithm(31). The assignment of phosphorylation sites was based on the PhosphoRS probability using the threshold of 75%. TMT reporter ions were extracted and normalized to the average of total phosphopeptide intensity detected in each channel. The phosphosite quantitation was calculated from the sum of the intensity of phosphopeptides with phosphorylation on the same residues.

### Quantitative SILAC-based phosphoproteomic analysis

#### Protein extraction, in-solution trypsin digestion, and desalting

A SILAC label incorporation efficiency check was performed for the three cell lines prior to the actual SILAC experiment to ascertain > 95% label incorporation efficiency. After cells were successfully labeled with the corresponding SILAC amino acids, the MCF10A parental- and two *PTEN* knockout cell lines were cultured in triplicates for protein harvest. 3 mg of protein from each cell line were combined to reach 9 mg of proteins per replicate. The same protocol was used for protein extraction, in-solution trypsin digestion, and peptide desalting for the TMT study as described above. The eluted peptide samples were lyophilized to dryness.

#### anti-pTyr-immunoprecipitation (IP)-based phosphopeptide enrichment

2 mg out of the total 9 mg peptides were used for pY enrichment. The immunoprecipitation procedure was carried out according to the manufacturer’s protocol (PTMScan® HS Phospho-Tyrosine (P-Tyr-1000) Kit, #38572, CST). Briefly, the lyophilized peptides were completely dissolved into the provided 1X HS immunoaffinity purification (IAP) bind buffer and adjusted for pH with 1 M Tris base buffer to neutral pH 7.0. The solution was cleared by centrifugation for 5 min at 10,000 x g at 4 °C. 20 µl of bead slurry was washed with ice-cold PBS 4 times for each IAP reaction. Peptide samples were incubated with bead slurry on an end-over-end rotator for 2 hours at 4 °C. Beads were collected on a magnetic separation rack and washed with chilled HS IAP Wash Buffer (1x) for a total of 4 times, followed by 2 times water washes. 50 µl of 0.15% TFA was used to elute the enriched peptides for a total of 2 times.

#### LC-MS/MS analysis

Phosphopeptides enriched by p-Tyr-1000 were analyzed on a reverse-phase LC system interfaced with an Orbitrap Exploris™ 480 Mass Spectrometer (Thermo Fisher Scientific) in DDA mode. 15 µl of the peptides reconstituted in 0.1% formic acid were injected into an analytical column (Acclaim PepMap 100, C18, 2 µm particle size) and separated with a 120 min gradient from 2% to 45% acetonitrile in 0.1% formic acid at a flow rate of 0.3 µl/min. The total run time was set to 140 min. Precursor MS scans were acquired in the range of 380–1,800 m/z with a resolution of 120,000 at 200 m/z on the Orbitrap analyser. The normalized AGC target for MS1 was set as 300% and ion filling time set 100 ms. The most intense ions with charge state ≥2 were isolated in 2 s cycle and fragmented using HCD fragmentation with 30% normalized collision energy and detected at a mass resolution of 30k at 200 m/z. The normalized AGC for MS/MS was set as 70%, and the ion filling time was set at 200 ms. Dynamic exclusion was set for 40 s with a 7 ppm mass window.

#### Mass spectrometry data analysis

Like the TMT study, Proteome Discoverer (v 3.0) was used for database search and p-Tyr-1000 enriched peptide identification and quantitation. The acquired MS/MS data were searched using SEQUEST search algorithms against the Human Uniport protein database (20,395 protein sequences) supplemented with frequently observed contaminants. The search parameters were set up as follows: (1) trypsin as a protease with full specificity; (2) a maximum of 2 allowed missed cleavages; (3) fixed modification: carbamidomethylation of cysteine; (4) variable modification: N-terminal acetylation, oxidation at methionine, phosphorylation at serine, threonine, and tyrosine; and SILAC labeling (^13^C_6_,^15^N_2_-lysine, 2H4-lysine, ^13^C_6_-arginine, and ^13^C_6_, ^15^N_4_-arginine). (5) MS tolerance was set at 10 ppm. (6) MS/MS tolerance was set to 0.02 Da. (7) The score cutoff value was set to 1% FDR at the peptide level. The probability of phosphorylation for each Ser/Thr/Tyr site on each peptide was calculated by the PhosphoRS algorithm(31). The assignment of phosphorylation sites was based on the PhosphoRS probability using a threshold of 75%. Since peptides with phosphotyrosine sites were enriched by pY1000, tyrosine residues were preferably assigned for the ambiguous phosphorylation sites. The abundance of all phosphopeptides in each sample was normalized to the average of total sum signal intensities in each sample.

The MS data were deposited to the ProteomeXchange Consortium via the PRIDE partner repository(32) and are available with the accession number PXD057520. Reviewers can access the dataset by logging in to the PRIDE website using the following account details:

Username

Password: juPz5t92uxVs

### Pathway and kinase enrichment and statistical analysis

For both TMT and SILAC studies, intensities of proteins, phosphosites, or phosphopeptides were averaged for the three biological replicates. Two *PTEN* knockout clones were compared with the parental MCF10A, respectively. The fold change cut-off was set as 1.5 and 0.67, while a two-sample Student’s T-test was performed (two-tailed distribution, two-sample equal variance), and p-value < 0.05 was set as the significance cut-off. Proteins or phosphosites that satisfied both thresholds for fold change and the p-value in two PTEN knockout clones were deemed as significantly differentially expressed. Missing values were replaced by 20% of the lowest value measured for each protein, phosphosite, or phosphopeptide. The kinase library with group/family information for each kinase was downloaded from Kinase.com (33). Gene Set Enrichment Analysis (GSEA) was done for total proteome in the TMT study using the default setups (version 4.3.2; permutation type: gene_set)(34). KEGG pathway enrichment analysis was performed on the DAVID platform with default parameters(35). Kinase enrichment analysis was conducted by KEA3 with default settings (36). The subcellular localization of human proteome was referenced from the Human Proteome Atlas (HPA) (37).

Breast cancer patient survival analysis was performed with the r packages: “survminer” and “survival”. Patient data were acquired via the Cancer Genome Atlas (TCGA) cBioPortal (Breast Invasive Carcinoma (TCGA, Provisional))(38). RNA expression data from DepMap https://depmap.org/portal/ (Expression Public 23Q2) were downloaded for 68 breast cell lines for the correlation analysis between PTEN and EphA2. Src interactome was downloaded from BioGRID (Version 4.4)(39), and interactors with significant fold changes in phosphorylation and expression were included (p value < 0.05; fold change >= 1.5). The substrates of Src were searched based on PhosphoSitePlus. The heatmaps were generated in R with the package “ComplexHeatmap”(40). Figures were mainly plotted in R with the ggplot2 package.

## Results

### Loss of PTEN promotes oncogenic phenotypes

We previously created a *PTEN* KO cell line model using MCF10A cells, a non-tumorigenic human breast epithelial cell line and found that *PTEN* deletion promoted EGF-independent cell proliferation, apoptotic resistance, and increased doxorubicin susceptibility (23). Given that MCF10A cells exhibit distinct characteristics when cultured in a medium with or without EGF, and the impact of PTEN loss is more pronounced under low EGF conditions(23), we examined the effect of culturing MCF10A cells and MCF10A PTEN-KO cells in 0.2 ng/ml EGF, which corresponds to 1% of the EGF typically present in normal MCF10A cell culture medium. Distinct morphological changes were observed in MCF10A cells and both clones with *PTEN* knockout **(Fig. 1A)**. In contrast to the typical cobblestone-like morphology of parental MCF10A cells, both *PTEN* KO clones exhibited larger cell size and an increased cytoplasm-to-nucleus ratio. *PTEN* KO cells also displayed a significant growth advantage compared to parental cells in the medium with 0.2 ng/ml EGF or the basal medium without EGF **(Fig. 1B)**, consistent with the previous finding that *PTEN* knockout permitted EGF-independent growth compared to parental cells. Functional validation of *PTEN* knockout was seen in the increased levels of phosphorylated AKT and ERK1/2 **(Fig. 1C)**.

**Figure 1.**
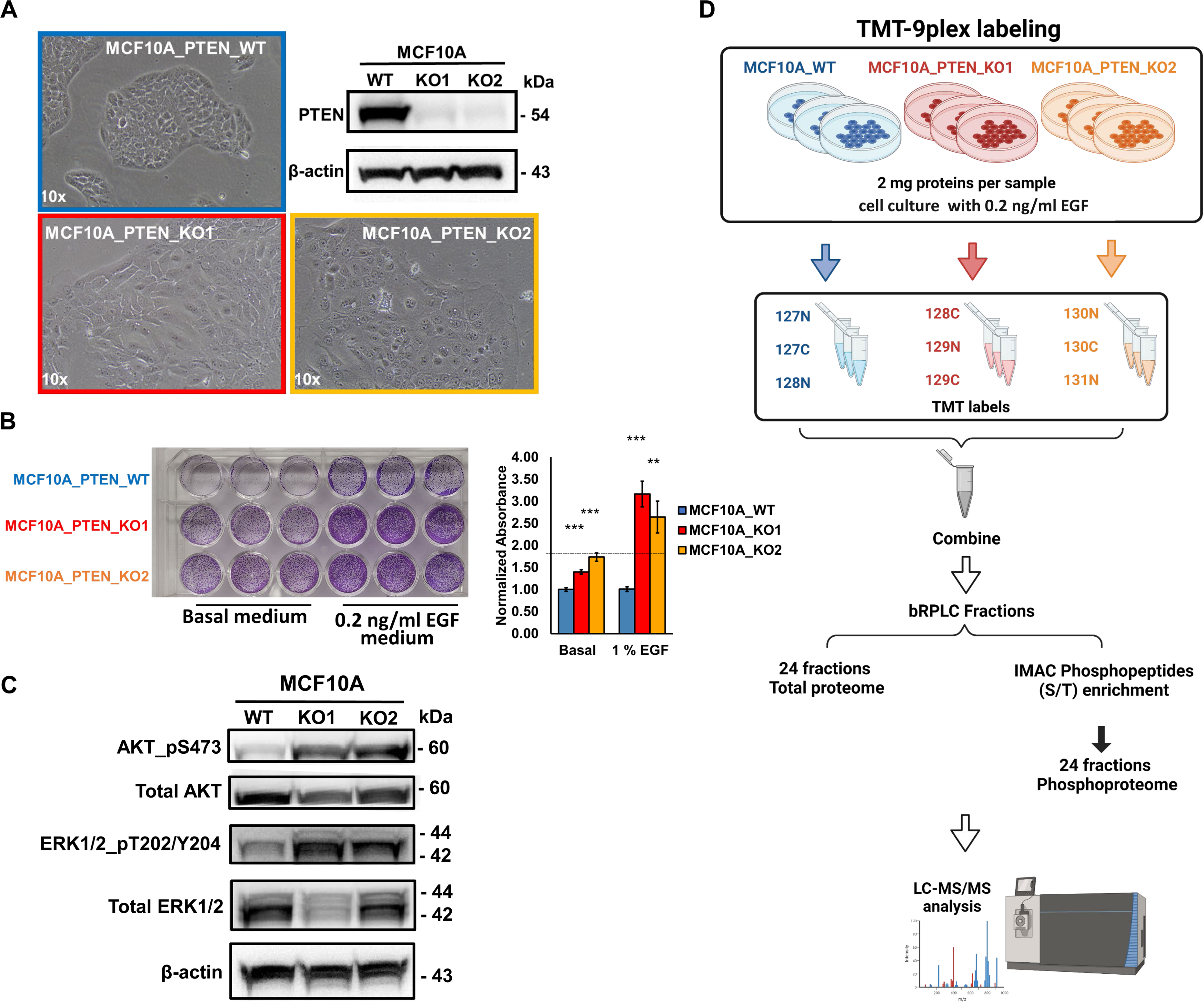
MCF10A *PTEN* knockout cell line models and proteomic and phosphoproteomic workflows. **(A)** The cell morphology of MCF10A cells with or without *PTEN* knockout (10x objective). Two MCF10A *PTEN* knockout cell clones (KO1 and KO2) were included in this study. Western blot confirmed the successful knockout of *PTEN*. MCF10A cells were cultured in 0.2 ng/ml EGF (1% of the EGF in MCF10A normal culture medium). WT: wide type; KO: knockout **(B)** Cell proliferation of MCF10A cells with or without *PTEN* knockout in basal or 1% EGF (0.2 ng/ml) cell culture medium. Cells were seeded in triplicates and cultured for 4 days prior to crystal violet staining. Stained crystal violet was eluted and quantified. **: p < 0.01; ***: p < 0.001 (Student’s T-test). **(C)** Western blot was used to check the expression levels of phosphorylated- and total AKT and p42/44 MAPK in *PTEN* wild-type or KO cells. **(D)** A schematic depiction of the proteomic study workflow. In this study, we started by carrying out TMT-9xplex labeling-based total proteomics and phosphoproteomics (IMAC enrichment) for MCF10A *PTEN* wild-type and KO cells.

### Quantitative analysis of the proteome and phosphoproteome of PTEN knockout cells

To comprehensively investigate the impact of *PTEN* knockout on other downstream targets, a TMT-labeling-based quantitative mass spectrometry analysis was performed for total proteome and phosphoproteome in MCF10A wildtype cells and two *PTEN* knockout clones **(Fig. 1D; Supplemental Data 1 & 2)**. We employed a 9-plex TMT labeling strategy with triplicates for each cell line. Deep profiling was achieved by fractionation of tryptic peptides into 24 fractions for both total proteome and phosphoproteome analysis **(Fig. 1D)**. 7,361 proteins from the total proteome and 9,627 phosphosites corresponding to 3,506 proteins from the phosphoproteome were identified with high confidence **(Fig. 2A)**. Among the identified phosphorylated sites, 84.7% (8,150 sites) were phosphorylated on serine, 14.7% (1,412 sites) were phosphorylated on threonine, and 0.68% (65 sites) were phosphorylated on tyrosine **(Fig. 2B)**. Principal component analysis (PCA) using the global proteomic- and phosphoproteomic data illustrated a clear separation between MCF10A cells and two PTEN KO clones, with tight clustering of replicate samples, indicating significant differences between MCF10A cells and the two *PTEN* knockout clones. It also underscored the rigor of our experimental procedures **(Fig. 2C-D)**. Of note, the two *PTEN* knockout clones were also comparably distinct from each other, suggesting that despite their shared origin of the same parental cell line and the common loss of PTEN, they exhibit distinct proteomic and phosphoproteomic profiles, albeit with similarities.

**Figure 2.**
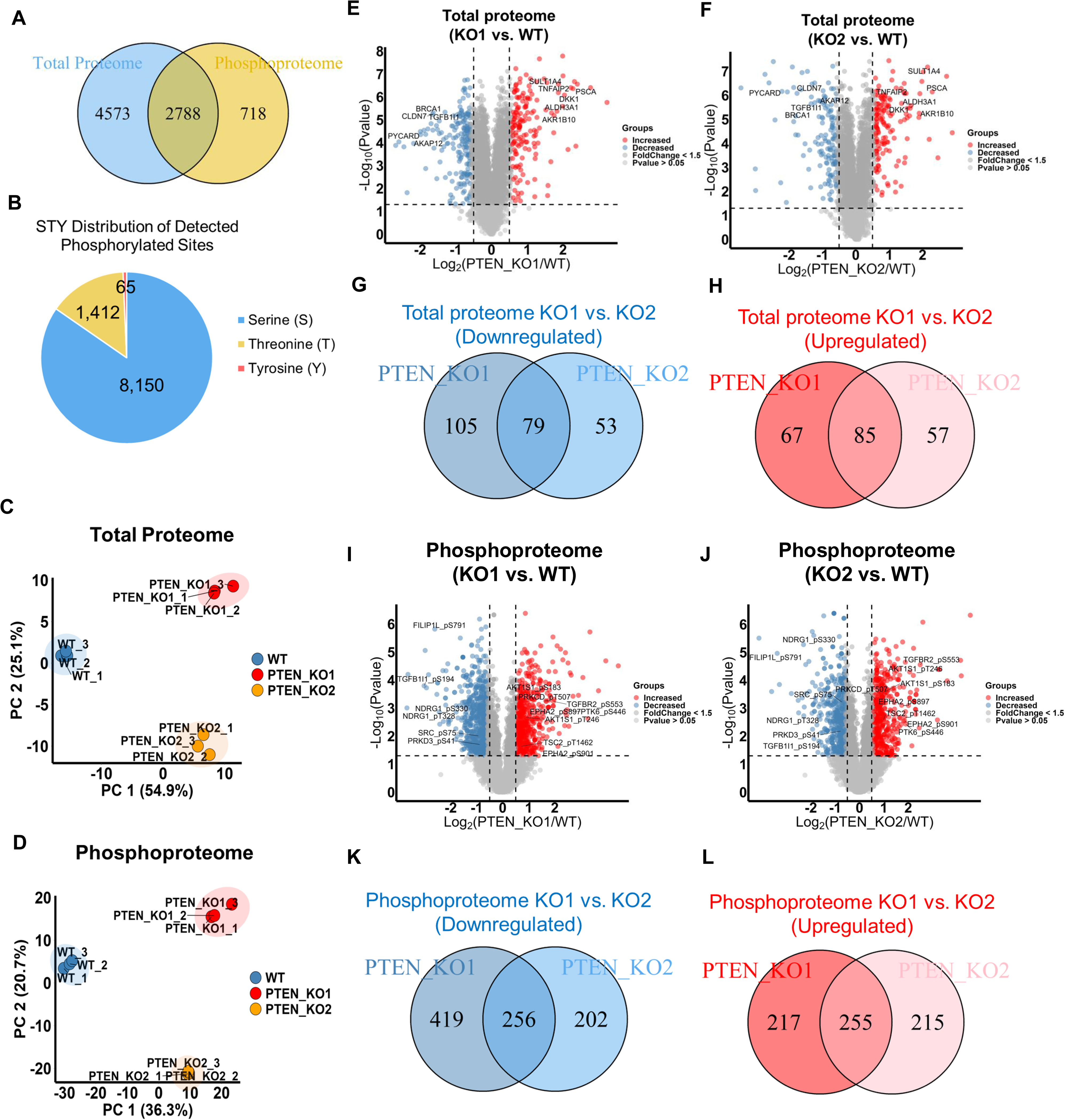
Quantitative proteomic and phosphoproteomic analysis for *PTEN* KO cell. **(A)** A Venn diagram showing the numbers of proteins identified and quantified in the total proteome and phosphoproteome. **(B)** A pie chart showing the numbers of phosphorylated serine, threonine, and tyrosine sites identified from the IMAC-enriched phosphoproteomic analysis. **(C-D)** Principal Component Analysis (PCA) plots of the total proteome and phosphoproteome. Proteins or phosphosites with high identification scores from PD and valid quantification were included in the analysis. **(E-F)** Volcano plots showing significantly altered proteins between MCF10A wide type and *PTEN* knockout clone 1 **(E)** or clone 2 **(F)** (Student’s T-test p value < 0.05). Red color dots indicate upregulated proteins after *PTEN* knockout (fold change >= 1.5), while blue dots indicate downregulated proteins after *PTEN* knockout (fold change <= 0.67). Proteins discussed in the main text were labeled. **(G-H)** Venn diagrams showing the overlapped proteins with significantly altered expression levels between the two MCF10A *PTEN* knockout clones. Significantly downregulated- **(G)** and upregulated **(H)** proteins were plotted separately (p value < 0.05; fold change >= 1.5). **(I-J)** Volcano plots showing the proteins with significantly altered phosphosites between MCF10A wide type and *PTEN* knockout clone 1 **(I)** or clone 2 **(J)** (Student’s T-test p value < 0.05). Red color dots indicate upregulated phosphosites after *PTEN* knockout (fold change >= 1.5), while blue dots indicate downregulated phosphosites after *PTEN* knockout (fold change <= 0.67). Proteins discussed in the main text were labeled. **(K-L)** Overlap of the significantly differentially expressed phosphosites between the two MCF10A *PTEN* knockout clones. Significantly downregulated- **(K)** and upregulated **(L)** phosphosites were plotted separately (Student’s T-test p value < 0.05; fold change >= 1.5).

Using a significance threshold of p> 0.05 and fold changes > 1.5 as cutoffs, we found that approximately 4.6% (336/7,361) **(Fig. 2E)** of the identified proteins and 12% (1,147/9,627) **(Fig. 2I)** of the identified phosphosites were significantly altered in MCF10A *PTEN* knockout cell clone 1, while 3.7% (274/7,361) **(Fig. 2F)** of the identified proteins and 9.6% (928/9,627) **(Fig. 2J)** of the identified phosphosites were significantly dysregulated in knockout cell clone 2, compared to MCF10A parental cells. Based on the presumption that the overlap of altered proteins between the two knockout clones would have the highest biological relevance, approximately 49% (164/336) of the altered proteins and 45% (511/1,147) of the altered phosphosites identified in knockout clone 1 were also shown in clone 2, while 60% (164/274) of the altered proteins and 55% (511/928) of the altered phosphosites in knockout clone 2 were significantly altered in clone 1 **(Fig. 2G-H, 2K-L)**. These overlapping dysregulations suggest shared molecular responses to PTEN loss in both clones, despite other differences in their proteomic and phosphoproteomic profiles.

### PTEN loss alters kinase and phosphatase signaling

In further analysis, we specifically examined the kinases detected in this study. The total proteomic and phosphoproteomic studies respectively identified 254 and 208 kinases (504 phosphorylated sites on the 208 kinases) covering all kinase categories, within which 163 kinases were identified in both studies **(Fig. 3A-B)**. Statistical analysis revealed that the protein expressions of CAMK2G, EphA2, MYLK, and TGFBR2 were significantly altered (p value < 0.05 & fold change >= 1.5) with only MYLK downregulated in *PTEN* knockout cells **(Fig. 3C & Supplemental Data 4)**. The top significantly upregulated phosphosites on the identified kinases were TGFBR2_pS553, PTK6_pS446, EphA2_pS901, CHEK2_pS260 and AKT3_pS34, while TTK_pS821, MAST4_pS206, ARAF_pS257, WEE1_pT190, PRKD3_pS41, and Src_pS75 were the top significantly downregulated sites on phosphorylated kinases **(Fig. 3D & Supplemental Data 4)**. Immunoblotting confirmed the changes in levels of protein kinases such as EphA2 and TGFBR2, as well as other signaling proteins such as SPHK1, DKK1 and RAB27B **(Fig. 3E-J)**.

**Figure 3.**
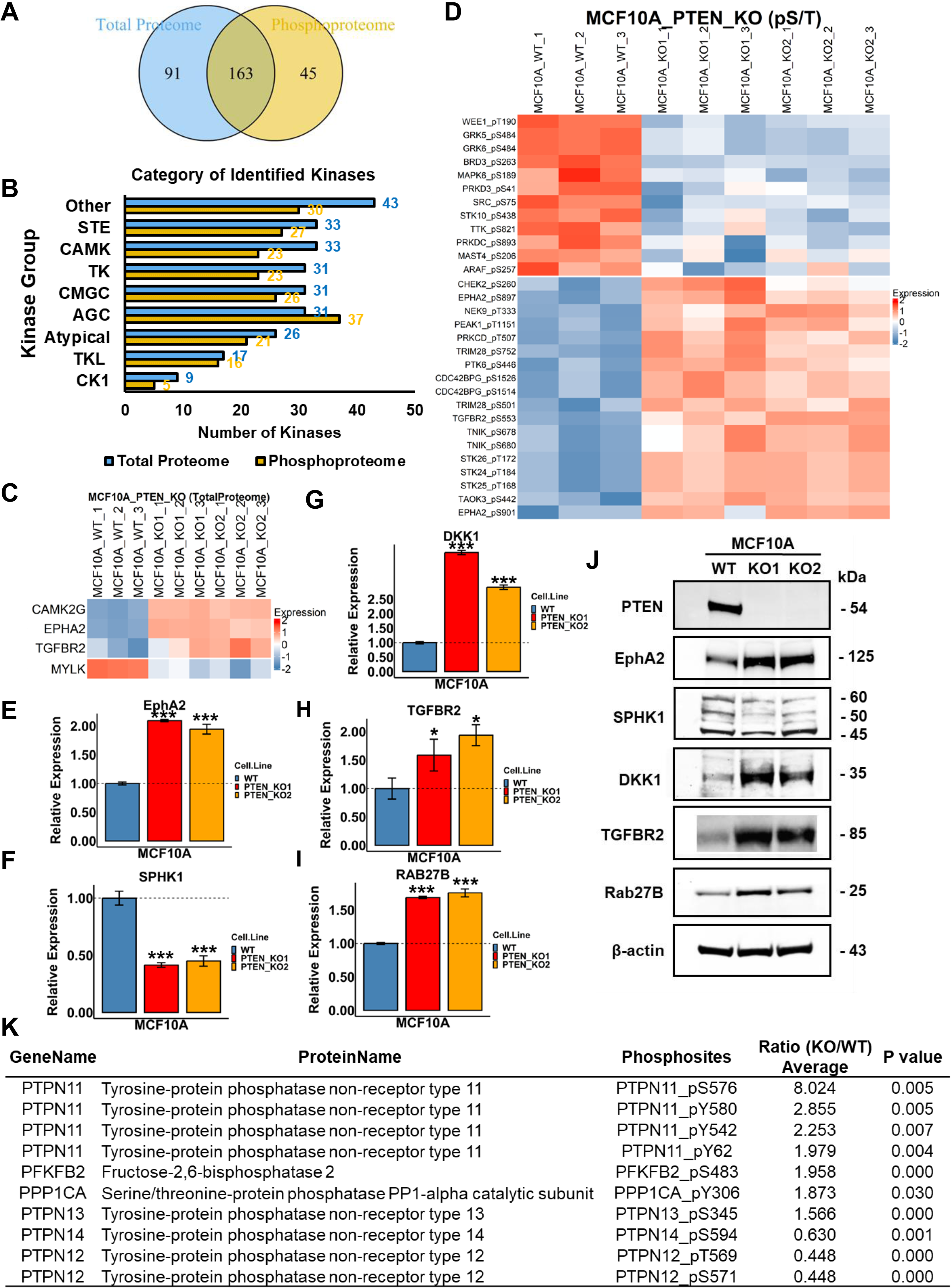
Kinases and phosphatases identified to be dysregulated in *PTEN* KO cells. **(A)** 254 and 208 kinases were identified in the total proteomic- and phosphoproteomic studies, respectively, within which, 163 kinases were found in both datasets. **(B)** Numbers of kinases identified in each category for the total proteomic- and phosphoproteomic datasets. STE: Homologs of yeast Sterile 7, Sterile 11, Sterile 20 kinases; CAMK: Calcium/calmodulin-dependent protein kinase; TK: Tyrosine kinase; CMGC: Containing CDK, MAPK, GSK3, CLK families; AGC: Containing PKA, PKG, PKC families; TKL: Tyrosine kinase–like; CK1: Casein kinase 1. **(C)** A heatmap showing the expression levels of four indicated kinases that were significantly altered in *PTEN* KO cells. **(D)** A heatmap showing the expression levels of the phosphorylation sites identified on kinases that were significantly up- or downregulated by *PTEN* KO. **(E-I)** Bar plots showing the relative expression levels of important signaling proteins detected in the proteomics analysis (as indicated) in *PTEN* KO clones compared to MCF10A parental cells. (Student’s T-test p value < 0.05: *, p value <0.01: ***); (**J**) The Immunoblot confirming the expression change of the indicated signaling proteins in MCF10A parental cells and PTEN KO cells. **(K)** A list of phosphorylated STY sites identified on phosphatases were significantly up- or downregulated by PTEN (Student’s T-test p value < 0.05; fold change >= 1.5). The kinase and phosphatase libraries used in this study were referenced from literature (33, 67).

Homeostasis of the cellular phosphoproteome is orchestrated by the seamless cooperation of kinases and phosphatases. Hence, we extended our analysis to phosphatases affected by PTEN. Though protein expression of phosphatases was generally unaltered **(Supplemental Data 4)**, phosphorylation of protein tyrosine phosphatases (SBF1-pT1138, PTPN11-pS576, pY580, pY542 and pY62, PTPN13-pS345, PTPN14-pS594, PTPN12-pT569 and pS571) and serine/threonine phosphatase PPP1CA at site pY306 were significantly dysregulated by PTEN loss (p value < 0.05 & fold change >= 1.5) **(Fig. 3K)**. Of note, the major phosphatases affected by PTEN were protein tyrosine phosphatases such as PTPN11 which has been extensively studied in tumor progression (41), highlighting the potential of PTEN in tyrosine phosphorylation.

### PTEN loss affects diverse cellular processes and signaling/metabolic pathways

To explore the biological activities affected by PTEN loss, we performed gene set enrichment analysis (GSEA) for the total proteome and KEGG pathway enrichment analysis specifically focusing on proteins exhibiting significantly altered phosphorylation levels **(Fig. 4)**. The observed negative enrichment of downregulated gene sets and positive enrichment of upregulated gene sets resulting from PTEN downregulation provided strong validation for the reliability of the proteomic dataset obtained in this study **(Fig. 4A)**. Protein expression of genes enriched in PTEN GSEA gene sets were plotted for MCF10A wild type and *PTEN* knockout clones **(Fig. S1)**. These PTEN-associated genes are involved in a collection of cell activities including cytoskeleton structure organization, metabolic reactions, extracellular exosome formation, and apoptosis. Apart from the signature gene sets associated with PTEN loss, we also observed that gene sets related to dysregulated key proteins within the PI3K-AKT signaling pathway, including mTOR and AKT pathways, were also enriched. These findings aligned well with our data (**Fig. 1**) and further support the activation of the PI3K-AKT pathway in the context of PTEN loss (42).

**Figure 4.**
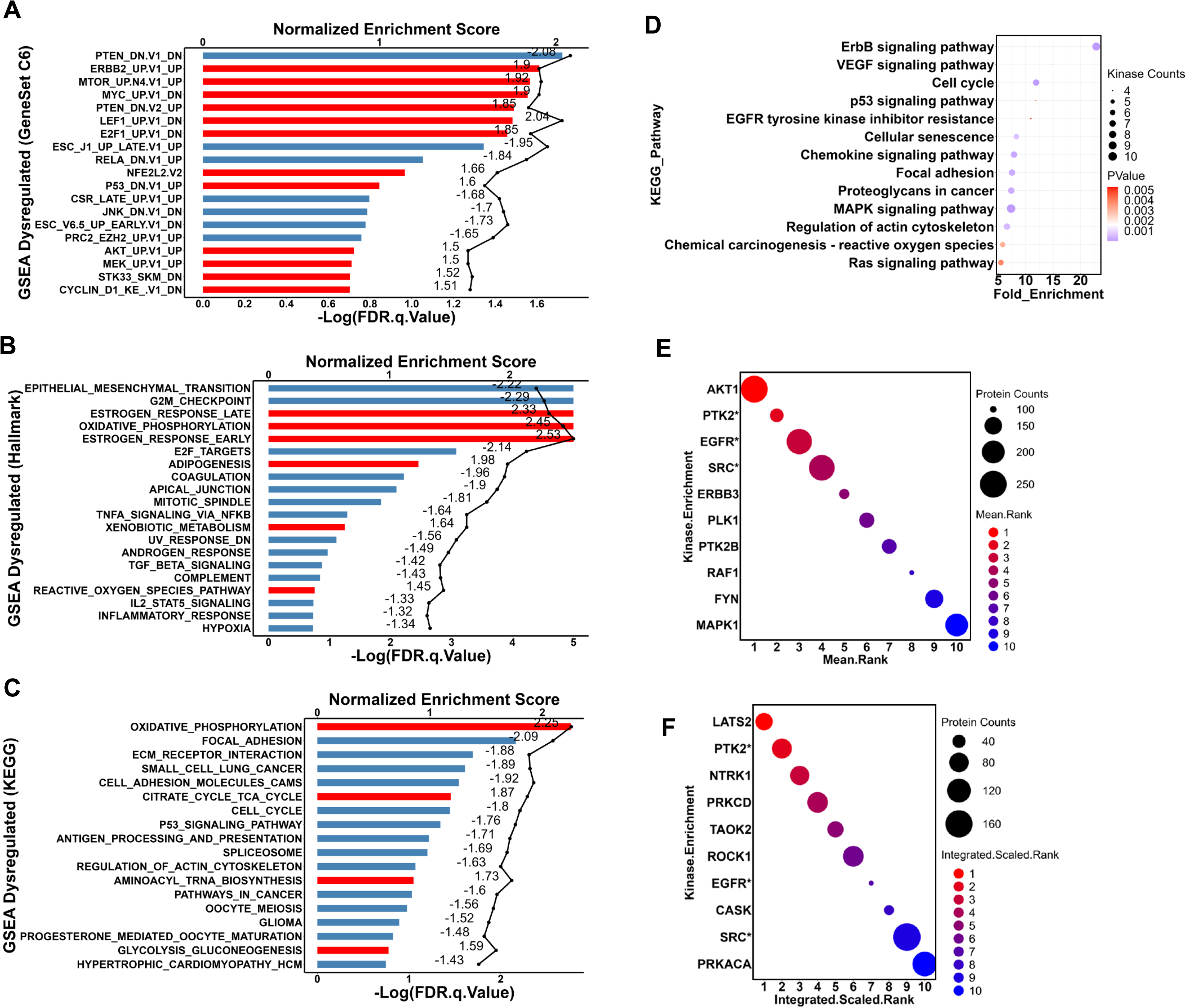
Pathway enrichment analysis. **(A-C)** Gene Set Enrichment Analysis (GSEA) of the proteins identified in the global proteomic analysis with significant (p<0.05) differential expression between MCF10A-WT and *PTEN* knockouts. Oncogenic signature- **(A)**, Hallmark- **(B)**, and KEGG- **(C)** gene sets were used for analysis. Bars represent FDR. Lines represent enrichment scores. The normalized enrichment scores were labeled on the graph. Red: positively enriched with positive scores; Blue: negatively enriched with negative scores. Enriched gene sets with FDR <= 0.1 for the Oncogenic gene sets, FDR <= 0.2 for the Hallmark gene sets, and FDR <= 0.2 for the KEGG gene sets were plotted. **(D)** Kinases with phosphorylation levels significantly altered (p < 0.05, consistent in both clones) were included in the KEGG pathway enrichment analysis via DAVID (Version Dec 2021). Default DAVID parameter setup was applied. **(E-F)** Proteins with phosphorylation levels significantly altered (p value < 0.05, fold change >= 1.5, consistent in both clones) were included in the kinase enrichment analysis in KEA3 with default setups. Results from both mean rank **(E)** and integrated scaled rank **(F)** analysis were plotted. X-axes represent the rank positions from 1 to 10. Kinases marked with * were identified in both methods.

Furthermore, an array of important oncogenic pathways was upregulated in *PTEN* KO cells, including ERBB, MYC, p53, JNK, MEK, and cyclin D1 pathways **(Fig. 4A)**. Apart from pivotal genes in signal transduction, gene sets of transcription factors (TF), namely MYC, E2F1, RELA (NF-kB), and NFE2L2, were also enriched, highlighting the impact of PTEN loss on transcriptional regulation. The Hallmark gene set enrichment analysis also uncovered a variety of cellular processes affected by PTEN, such as epithelial-mesenchymal transition (EMT), cell cycle, metabolism, oncogenesis, and immune responses **(Fig. 4B)**. Further, KEGG pathway analysis revealed that central carbon metabolism was activated by PTEN loss, as evidenced by the positive enrichment of oxidative phosphorylation, citrate cycle, and glycolysis metabolic pathways **(Fig. 4C and Fig. S2)**.

We further conducted KEGG pathway enrichment of non-phosphorylated and phosphorylated proteins that were significantly changed by PTEN knockout (p value < 0.05, fold change >= 1.5) using DAVID (35) **(Fig. S3A-B & Supplemental Data 5)**. This analysis showed that the PI3K-AKT signaling pathway is activated by PTEN loss, as reflected by changes in proteins and phosphosites, such as BRCA1, EphA2, Ephrin A1, integrins, and TSC2 (**Fig. S3C-Q)**. Having identified the phosphorylation sites of kinases deregulated by PTEN loss **(Supplemental Data 4)**, we conducted KEGG pathway enrichment analysis of these kinases to gain insights into the kinase-enriched signaling pathways **(Fig. 4D)**. A variety of cancer-related signaling pathways were revealed to be associated with PTEN loss, namely, ERBB, VEGF, p53, EGFR, MAPK, reactive oxygen species (ROS) and Ras. Moreover, cell cycle and cellular senescence, along with focal adhesion and actin cytoskeleton signaling pathways, were also enriched in PTEN-knockout cells.

To identify potential upstream kinases activated by PTEN loss, we performed kinase enrichment analysis of the significantly changed phosphoproteins using the Kinase Enrichment Analysis 3 (KEA3) algorithm (36) and identified the top ten enriched kinases using two integrated methods, respectively **(Fig. 4E-F & Supplemental Data 5)**. Notably, the serine-threonine protein kinase AKT1 was ranked the top kinase by the mean rank method. Additionally, tyrosine kinases, PTK2, EGFR, and Src consistently appeared among the top ten enriched kinases using both methods. Of significance, these findings, along with our GSEA analysis, suggested that protein tyrosine kinases, in addition to serine/threonine kinases, may also play significant roles in the context of PTEN loss and could potentially be activated in response to PTEN inactivation.

### PTEN loss regulates global tyrosine phosphorylation and tyrosine kinases

Our analysis indicating that PTEN loss may potentially enhance tyrosine kinase-mediated signaling led us to assess the effects of *PTEN* knockout on tyrosine kinases. To survey the broad effects of PTEN loss on tyrosine kinases, we used an anti-pan phosphotyrosine antibody, 4G10, to examine tyrosine phosphorylation levels in MCF10A wild-type and *PTEN* knockout cells and MCF10A cells with or without siRNA mediated PTEN knockdown. These studies indicated that PTEN suppression by either knockout or transient knockdown substantially increases tyrosine phosphorylation **(Fig. 5A).** We then carried out a three-state SILAC labeling-based quantitative phosphoproteomic study to quantify phosphorylated tyrosine peptides. After enrichment by immunoaffinity precipitation (IAP) with pY-1000 antibodies(43) **(Fig. 5B-M & Supplemental Data 6 & 7),** we identified 2,080 tyrosine-phosphorylated peptides. Notably, there was a high degree of overlap between each replicate study, with over 93% (1,944/2,080) of the phosphorylated peptides being identified in all three replicate studies **(Fig. 5C)**. We further analyzed the phosphotyrosine data on the phosphosite level. Consistent with what we observed in our total proteomics and IMAC-based phosphoproteomics study, the MCF10A wild-type cells and two *PTEN* knockout clones demonstrated distinctive phosphotyrosine profiles **(Fig. 5D**). In contrast to the relatively balanced number of proteins and phosphosites with upregulation or downregulation observed in the global proteome and IMAC-enriched phosphoproteome (**Fig. 2E-F** and **I-J**), there was a significantly greater number of upregulated tyrosine phosphorylation sites compared to downregulated ones in both *PTEN* KO clones, compared to MCF10A cells. Specifically, there were 311 hyperphosphorylated pTyr sites in PTEN-KO1 and 236 in PTEN-KO2 cells, while only 35 were hypophosphorylated in PTEN-KO1 and 32 in PTEN-KO2 cells **(Fig. 5E-H)**. Considering the phosphorylation sites that were consistently altered in both clones, among the 663 identified and quantifiable sites, 201 pY sites (30%) were significantly altered (p-value < 0.05, fold change >= 1.5), with the majority (89%, 179 pTyr sites) being upregulated **(Supplemental Data 6)**. These data indicated that protein tyrosine phosphorylation levels were globally elevated to a substantial degree in cells with PTEN loss.

**Figure 5.**
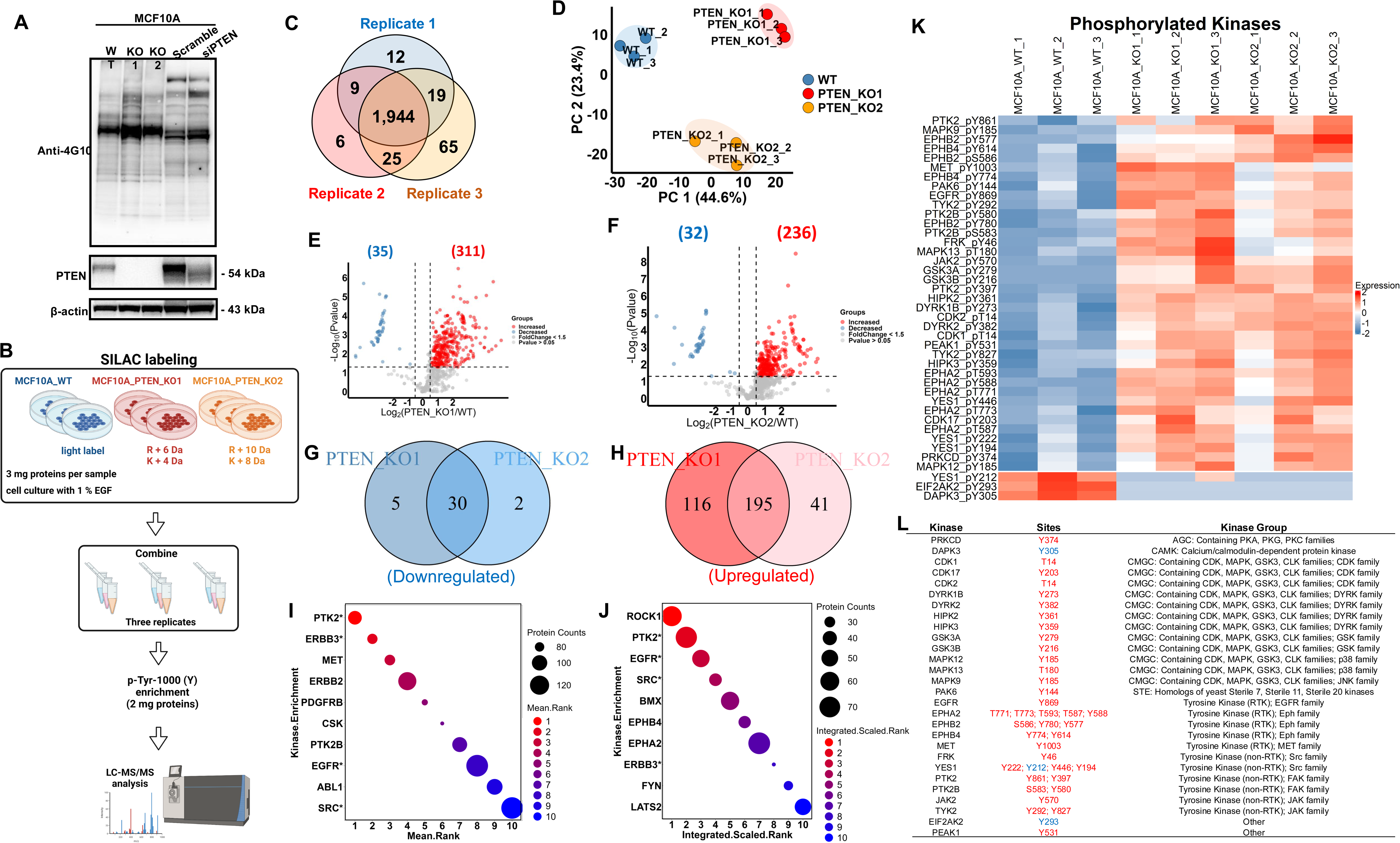
PTEN loss activated protein tyrosine phosphorylation and tyrosine kinases. **(A)** The global protein phosphotyrosine (pTyr) level in MCF10A-WT and *PTEN* knockouts and MCF10A cells with transient knockdown of PTEN siRNA or scramble siRNA. The pTyr expression levels were detected by Western blot using 4G10 anti-pan pTyr antibody **(B)** A schema showing SILAC labeling-based phosphoproteomic analysis of pTyr peptides enriched by pY-1000 antibodies from MCF10A cells, PTEN KO1 and KO2 cell lines. **(C)** A Venn diagram shows that over 93% (1,944 pTyr sites) of the identified pTyr sites are commonly identified among all three replicates. **(D)** PCA analysis of the pTyr data suggested distinct pTyr patterns among MCF10A cells and PTEN KO cells. **(E-F)** Volcano plots showed protein pTyr sites with significant expression levels between MCF10A wide type and *PTEN* KO clone 1 **(E)** or clone 2 **(F)** (Student’s T-test p value < 0.05). Red dots indicate upregulated phosphorylated sites after *PTEN* knockout (fold change >= 1.5), while blue dots indicate downregulated phosphorylated sites after *PTEN* knockout (fold change >= 1.5). **(G-H)** Venn diagrams depicted the pTyr sites commonly downregulated (**G**) or upregulated (**H**) in both PTEN KO clones compared to MCF10A cells. **(I-J)** Proteins with pTyr levels significantly altered (p value < 0.05, fold change >= 1.5, consistent in both clones) were included in the kinase enrichment analysis in KEA3 with default setups. Results from both mean rank **(I)** and integrated scaled rank **(J)** analysis were plotted. X-axes represent the rank positions from 1 to 10. Kinases marked with * were identified in both methods. **(K)** A heatmap showing significantly altered phosphorylation sites (mainly tyrosine sites) of protein kinases identified in the pTyr proteomic analysis. **(L)** A list of protein kinases with altered tyrosine phosphorylation in Figure 5K. Phosphorylation sites downregulated were marked in blue, while upregulated phosphorylation sites were marked in red.

To identify upstream tyrosine kinases potentially responsible for the tyrosine phosphorylation changes in *PTEN* KO cells, we adopted a similar approach that was used to analyze the IMAC-enriched phosphoproteomic data and performed kinase enrichment analysis of the significantly changed phosphoproteins using the KEA3 algorithm (36) for the phosphotyrosine dataset. As shown in Figures 5I and 5J **(Supplemental Data 7)**, PTK2, ERBB3, EGFR, and Src were among the top ten enriched kinases based on both analysis methods. Consistent with these results, the examination of the altered phosphopeptidome after *PTEN* knockout revealed the highest change in phosphorylation of proteins downstream of Src and EGFR, evidenced by the mean rank method **(Fig. 5J)**. Next, we specifically delved into kinases with altered tyrosine phosphorylation after *PTEN* knockout **(Fig. 5K-L & Supplemental Data 7)**. Five RTKs, EGFR, EphA2, EphB2, EphB4, and MET, and six non-RTKs including FRK, YES1, PTK2, PTK2B, JAK2 and TYK2, were identified to be associated with PTEN loss. Specifically, phosphorylation of several members from the Eph RTK family (EphB2_pY780 & pY577; EphB4_pY774 & pY614; EphA2_pY588) and the dual-specificity protein kinase family (DYRK1B_pY273; DYRK2_pY382; HIPK2_pY361; HIPK3_pY359) were affected by PTEN loss. Second to EphA2, which had 5 sites altered by PTEN loss, from the Src family Yes1 had 4 sites (Y222 & Y212 & Y446 & Y194) that were markedly affected, indicating the possibility that PTEN loss may exert its greatest impacts on various RTKs and non-RTKs. To systematically assess the impact of PTEN loss on kinase expression and phosphorylation, we mapped the significantly altered kinases (p-value < 0.05; both protein expression and phosphorylation) onto the kinase phylogenetic tree (33) **(Fig. 6A & Supplemental Data 8)**. The results showed that PTEN loss broadly influenced kinases across all categories, affecting both their expression and phosphorylation levels.

**Figure 6.**
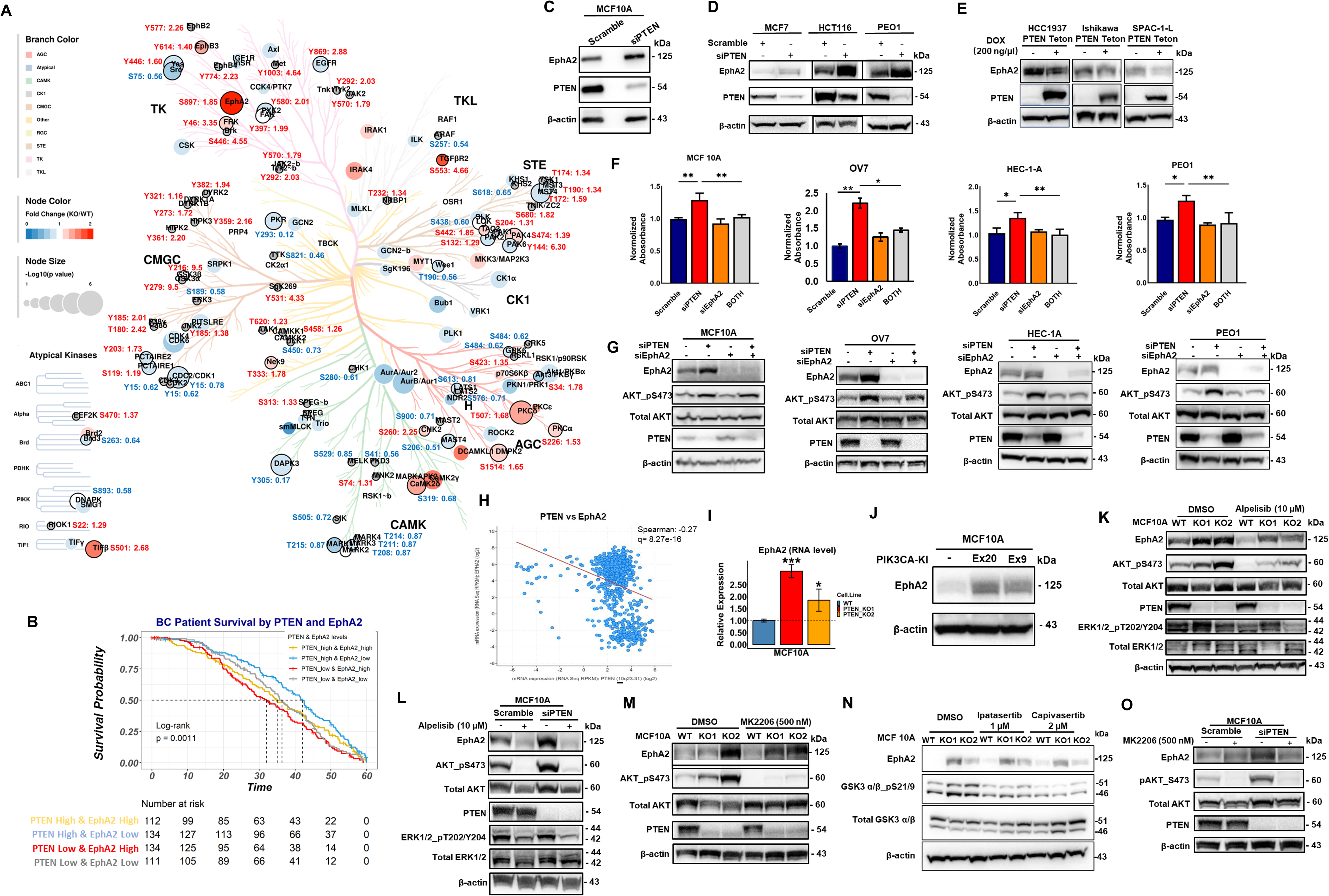
PTEN loss upregulates EphA2. **(A)** A phylogenetic tree of protein kinases whose phosphorylation levels and/or expression levels were significantly altered by *PTEN* knockout. The plot was generated using http://phanstiel-lab.med.unc.edu/CORAL/. Solid red circles: kinases upregulated in PTEN KO cells; Solid blue circles: kinases downregulated in PTEN KO cells; Grey circles: kinases detected but not significantly changed in expression; Circle sizes represent the p-value; Black circle strokes: kinases with significant changes in phosphorylation levels. Representative phosphorylation site for each kinase was labeled with site position and fold change (red: upregulation after PTEN loss; blue: downregulation after PTEN loss). **(B)** Kaplan Meier survival analysis of breast cancer patients with different expression levels of PTEN and EphA2. Breast cancer patient data were downloaded from the Cancer Genome Atlas (TCGA) cBioPortal (Breast Invasive Carcinoma (TCGA, Provisional))(38). p-value was calculated using the log-rank method. **(C-D)** Western blot analysis examined EphA2 expression level in MCF10A cells (**C**) and HCT116 and PEO1 cells (**D**) with siRNA-mediated PTEN transient knockdown. **(E)** PTEN expression was induced in PTEN deficient HCC1937, Ishikawa and SPAC-1-L cells. The expression levels of EphA2 and PTEN were examined by western blot. β-Actin serves as a loading control. **(F)** The expression levels of PTEN, EphA2, pAKT and AKT were examined by Western blot in MCF10A, OV7, HEC-1-A and PEO1 cells with siRNA knockdown of PTEN and/or EphA2 expression. (**G**) Crystal violet staining to evaluate the proliferation of MCF10A, OV7, HEC-1-A and PEO1 cells treated with scramble siRNA, siPTEN, siEphA2, and a combination of siPTEN and siEphA2. Quantification of the absorbance was plotted. (**H)** Correlation analysis of the mRNA expression of PTEN and EphA2 in 921 cancer cell lines was performed using Cancer Cell Line Encyclopedia dataset in cBioPortal. The correlation between PTEN and EphA2 was negative with Pearson correlation being −0.27 (p value = 8.27e-16). **(I)** RT-qPCR analysis of the mRNA levels of EphA2 in MCF10A wide type and *PTEN* knockout cells. **(J)** Knock-in (KI) of two gain-of-function mutations of *PIK3CA* (Ex20 and Ex9)(24) increased EphA2 expression. **(K-L)** EphA2 protein level was examined in MCF10A wide type and *PTEN* KO cell clones treated with PI3K inhibitor Alpelisib (10 µM) and MCF10A cells with (**K**) or without siPTEN knockdown (**L**). **(M-N)** EphA2 expression was examined in MCF10A, PTEN-KO clones treated with AKT inhibitors, MK2206 (**M**), ipatasertib or capivasertib (**N**). AKT pS473 and GSK3 α/β pS21/19 were used to evaluate the AKT inhibition by indicated AKT inhibitors. (O) EphA2 expression level was examined in MCF10A cells with siPTEN knockdown treated with MK2206.

### EphA2 is upregulated after *PTEN* knockout in an AKT-independent manner

The extensive impact of PTEN loss on tyrosine phosphorylation is likely mediated through the activation of RTKs and non-RTKs. As an RTK, EphA2 plays critical roles in cellular functions and is recognized as a promising therapeutic target in multiple cancers, including breast cancer (44, 45). Given the high frequency of genetic alterations in the PI3K signaling pathway, particularly involving *PIK3CA* and *PTEN*, in breast cancer (4), we investigated the relationship between patient survival and expression levels of PTEN and EphA2, using the Breast Invasive Carcinoma database (TCGA, Provisional) from cBioPortal (38). Patients were categorized into four groups based on the mRNA levels of PTEN and EphA2 in their tumors. Among these groups, the cohort with low PTEN expression and high EphA2 expression exhibited the poorest survival outcomes (**Fig. 6B**).

Our data consistently demonstrated that EphA2 expression and phosphorylation levels were significantly upregulated in PTEN knockout cells **(Fig. 3E and J & Fig. S4A)**. To confirm that this upregulation of EphA2 is specifically associated with PTEN loss rather than being a result of cellular adaptation during the process of establishing the *PTEN* knockout cell lines, we employed PTEN-specific siRNA to transiently knockdown PTEN expression in MCF10A cells. The results demonstrated that transient suppression of PTEN expression also led to a substantial increase in the expression of EphA2 **(Fig. 6C & Fig. S4B)**, supporting the direct association between PTEN loss and EphA2 upregulation. To validate that the upregulation of EphA2 by PTEN loss is not restricted to a single cell line or breast cancer, we conducted PTEN siRNA knockdown in two different cancer cell lines: HCT116 (colorectal cancer) and PEO1 (ovarian cancer). Remarkably, EphA2 expression levels were consistently upregulated across all three cancer cell lines following PTEN knockdown (**Fig. 6D**). Additionally, we generated inducible overexpression of PTEN in PTEN-deficient cell lines, including SPAC-1-L and Ishikawa (endometrial cancer lines) as well as HCC1937 (breast cancer) to demonstrate that EphA2 was downregulated in cancer cells with PTEN overexpression **(Fig. 6E)**. These results reveal a consistent inverse relationship between PTEN and EphA2 expression across multiple cancer types, highlighting the broader significance of the PTEN-EphA2 regulatory axis across diverse cancer contexts.

To investigate the functional role of EphA2 upregulation in PTEN-regulated signaling, PTEN and EphA2 were knocked down simultaneously in MCF10A, as well as ovarian cancer cells (OV7 and PEO1) and endometrial cancer cells (HEC-1-A). The enhanced proliferation induced by PTEN knockdown was abrogated by simultaneous knockdown of EphA2, suggesting that the increased proliferation rate of PTEN deficient cells depends at least partially on an increase in EphA2 (**Fig. 6F-G**). However, the knockdown of EphA2 alone had minimal impact on cell proliferation. These findings, together with the survival analysis (**Fig. 6B**), suggest that activation of EphA2 signaling contributes to cancer progression in the context of PTEN loss.

To investigate whether EphA2 upregulation upon PTEN suppression occurs at the transcriptional or translational level, correlation analysis was performed using mRNA expression data from the 921 cancer cell lines in the Cancer Cell Line Encyclopedia(46). A significant negative correlation was observed between PTEN and EphA2 mRNA expression levels (Spearman’s correlation: −0.268, q value: 8.27e-16) **(Fig. 6H)**. In line with this analysis, our real-time RT-PCR studies further confirmed that EphA2 mRNA expression levels were also higher in *PTEN* knockout clones, compared with the MCF10A parental cells **(Fig. 6I)**. These findings collectively demonstrate that EphA2 upregulation in response to PTEN loss mainly occurs at the transcriptional level.

Because PTEN acts as a negative regulator of the PI3K pathway, loss of PTEN function often leads to increased PIP3 levels and hyperactivation of the PI3K-AKT pathway(1). To further investigate the signaling regulation of EphA2 by PTEN, we examined EphA2 expression in our previously studied MCF10A cells with isogenic knock-in of oncogenic PIK3CA mutations (E545K and H1047R)(24). Activation of PI3K by oncogenic PIK3CA mutations also increased EphA2 expression **(Fig. 6J)**. Further supporting the role of PI3K in EphA2 regulation, treatment with the PIK3CA inhibitor alpelisib substantially reduced EphA2 protein expression in MCF10A wild-type cells and in two PTEN knockout clones **(Fig. 6K)**. Similar results were observed in MCF10A cells with siRNA-mediated PTEN knockdown (**Fig. 6L**) and in PEO1 cells treated with alpelisib (**Fig. S5A**). However, a substantial level of EphA2 expression persisted even after complete abrogation of AKT phosphorylation by alpelisib **(Fig. 6K and Fig. S5A)**, raising the possibility that additional mechanisms beyond PI3K may contribute to EphA2 expression in the context of PTEN loss.

Given that AKT and mTOR are key kinases downstream of PI3K, we investigated whether their activation is necessary for the EphA2 upregulation. Surprisingly, treatment of MCF10A wild-type cells, PTEN knockout cells, PTEN knockdown cells with three different AKT inhibitors (MK2206, ipatasertib, and capivasertib), effectively reduced AKT phosphorylation and/or downstream GSK3-α/β phosphorylation but did not result in consistent or significant EphA2 downregulation (**Fig. 6M-O & Fig. S5B**). Similarly, inhibiting mTOR with Rapamycin did not decrease EphA2 expression in either MCF10A (wild-type or PTEN knockout) or PEO1 cells (**Fig. S5C-D**). These results indicated that neither AKT nor mTOR serves as a primary regulator of EphA2 upregulation in the context of PTEN loss.

### EphA2 is regulated by the Src signaling pathway

Our quantitative proteomics and phosphoproteomics analyses demonstrated that PTEN loss significantly activates tyrosine kinase signaling cascades in addition to the canonical PI3K-AKT pathway. Kinase enrichment analyses (**Fig. 4E-F & Fig. 5I-J**) identified hyperactivation of non-receptor tyrosine kinases (non-RTKs) such as FAK and Src in PTEN-deficient cells. To examine their roles in EphA2 regulation, we treated cells with the FAK inhibitor GSK2256098 but found minimal effects on EphA2 expression in either parental MCF10A or PTEN knockout cells (**Fig. S5E**). Further analysis of our phosphoproteomics data using the Src interactome dataset from the BioGRID database (47), revealed extensive dysregulation in proteins interacting with or phosphorylated by Src following PTEN loss **(Fig. 7A & Supplemental Data 7),** suggesting that Src may be key in regulating EphA2 expression in PTEN-deficient cells.

**Figure 7.**
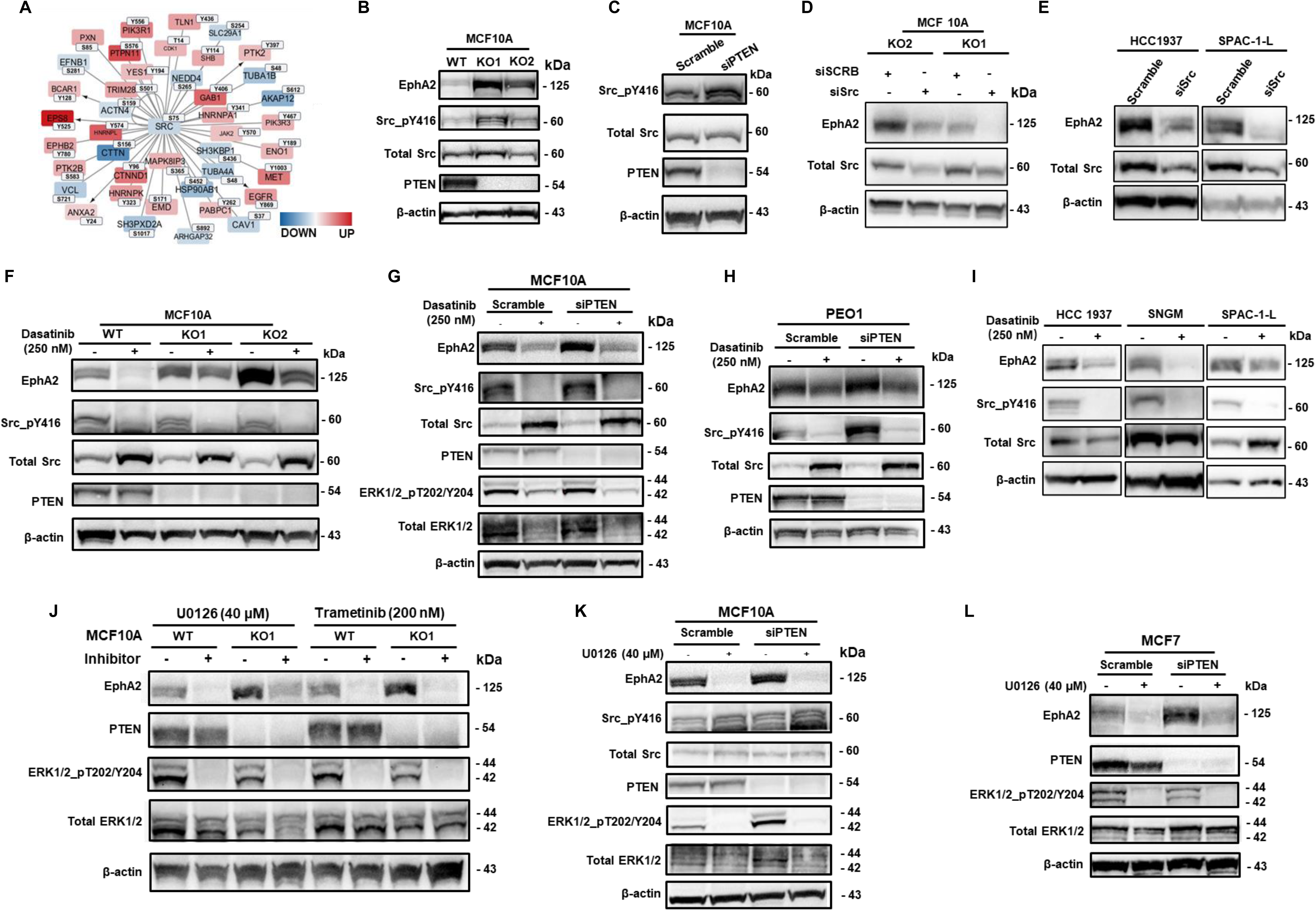
Src-MEK signaling mediates the upregulation of EphA2 in cells with PTEN loss. **(A)** Differentially regulated Src interacting phosphoproteins were identified using BioGRID(39) database (version 4.4), and the protein-protein interaction map was plotted using Cytoscape. Phosphorylated protein interactors marked in red were significantly upregulated in PTEN KO cells, while phosphorylated protein interactors marked in blue were significantly downregulated. The darker the color, the bigger the difference. The node size corresponded to p value, and bigger text size indicated smaller p value. One representative phosphorylation site of each protein was labeled for the corresponding proteins. The full list of differentially regulated phosphosites is shown in **Supplemental Data 7**. The edges with arrows indicate that these proteins are the substrates of Src. **(B-C)** PTEN knockout (**B**) and transient knockdown (**C**) in MCF10A cells induced Src-Y416 hyperphosphorylation. **(D-E)** EphA2 expression levels were measured in PTEN deficient MCF10A-PTEN-KD cells (**D**), HCC1937 and SPAC-1-L cells (**E**) with siRNA knockdown of Src. (**F-H**) Src inhibitor Dasatinib was used to treat MCF10A cells with *PTEN* knockout **(F)**, MCF10A cells (**G**) and PEO1 cells with PTEN transient knockdown (**H**). Src pY416 was used to evaluate the dasatinib inhibition efficiency. (**I**) EphA2 expression was checked by western blot in PTEN deficient HCC1937, SNGM and SPAC-1-L cells treated with dasatinib. **(J-L)** EphA2 expression was reduced by MEK inhibitors U0126 and Trametinib in MCF10A cells with *PTEN* knockout **(J)**, in MCF10A cells (**K**) and MCF7 cells (**L**) with transient siPTEN knockdown. Antibody targeting phosphorylated ERK1/2 at T202/Y204 was employed to evaluate the activation of MEK and the successful inhibition of MEK by kinase inhibitors U0126 and Trametinib.

To investigate the functional role of Src in cells with *PTEN* loss, we first examined the Src phosphorylation at tyrosine residue 416 (pY416) upon *PTEN* knockout. pY416 is a specific autophosphorylation site on the Src kinase activation loop, and the Y416 phosphorylation leads to Src conformational change, which is required for full kinase action (48). In line with our proteomics data analysis, Src was hyperphosphorylated and activated regardless of long-term *PTEN* knockout or transient PTEN knockdown in MCF10A cells **(Fig. 7B-C)**. To examine whether the Src kinase is involved in regulation of EphA2 expression, we used Src-specific siRNA to knock Src down in PTEN-deficient cells, including MCF10A-KO1 and -KO2 cells, HCC1937 (breast cancer) and SPAC-1-L (endometrial cancer) cancer cells and confirmed that suppressing Src expression can substantially reduce EphA2 expression across all tested cell lines (**Fig. 7D-E).** Previous studies have shown that dasatinib, an FDA-approved Src kinase inhibitor, can inhibit EphA2 autophosphorylation and protein expression in a dose-dependent manner (49). Using dasatinib, we could reduce EphA2 protein expression in MCF10A *PTEN* KO cell lines **(Fig. 7F)**, as well as in MCF10A cells with siRNA-mediated PTEN knockdown **(Fig. 7G)**. Likewise, in PEO1 ovarian cancer cells, PTEN knockdown activated Src kinase and increase EphA2 expression, whereas inhibiting Src with dasatinib potently suppressed Src phosphorylation and reduced EphA2 expression **(Fig. 7H)**. Similar results were observed in PTEN-deficient breast (HCC1937) and endometrial (SNGM and SPAC-1-L) cancer cells treated with dasatinib (**Fig. 7I**).

One of the key signaling pathways downstream of Src is the Ras-Raf-MEK-ERK cascades, wherein Src activates Ras, leading to phosphorylation of Raf, MEK, and ultimately ERK1/2 (50). As a Src inhibitor, dasatinib has been shown to inhibit ERK1/2 activation in a cell type-dependent manner(51, 52). In the present study, our GSEA analysis revealed that a MEK-activated signature and the MAPK signaling pathway were positively enriched in the *PTEN* knockout cells **(Fig. 4A & 4D)**. These findings suggest that MEK-ERK1/2 signaling likely acts downstream of PI3K and PTEN in MCF10A cells. Notably, EphA2 has been reported to be regulated by MEK(53, 54), reinforcing the potential role of MEK-ERK1/2 in EphA2 regulation. Our studies confirmed that inhibiting Src by dasatinib dramatically attenuated ERK1/2 phosphorylation in MCF10A cells **(Fig. 7G)**. Similarly, alpelisib also decreased phospho-ERK1/2 levels in MCF10A cells **(Fig. 6K-L)**. To confirm MEK’s role in EphA2 regulation, we treated MCF10A cells with the MEK inhibitors U0126 and trametinib, each of which significantly reduced EphA2 expression in wild-type and PTEN knockout cells **(Fig. 7J & Fig. S5F)**. Similar results were observed in MCF10A and MCF7 cells treated with U0126 after transient PTEN knockdown (**Fig. 7K-L**). The role of MEK in the regulation of EphA2 was also confirmed in another ovarian cancer cell line, PEO1 cells treated with U0126 **(Fig. S5G)**. These findings indicate that Src activation, rather than AKT, plays a critical role in upregulating EphA2 expression in PTEN-deficient cells.

### AKT inhibition and Src inhibition synergistically suppress cells with PTEN deficiency

Our findings demonstrate that PTEN loss activates both the canonical PI3K-AKT pathway and tyrosine kinase signaling, revealing a complex signaling network in PTEN-deficient cells that contributes to tumor progression. These results suggest that targeting AKT alone may be insufficient for optimal therapeutic efficacy in cancers with PTEN loss. Consequently, Src, EphA2, and MEK-ERK1/2 could emerge as promising additional targets for therapeutic intervention alongside AKT signaling.

To investigate the combinatorial effects of targeting both AKT and Src signaling in PTEN-deficient cancer cells, we treated MCF10A-PTEN-KO, breast cancer HCC1937 cells and endometrial cancer SPAC-1-L cells (all PTEN-deficient) with different concentrations of the Src inhibitor dasatinib (from 0 to 120 nM) and AKT inhibitor capivasertib (from 0 to 100uM), assessed proliferation, and performed synergy analysis. The synergy plots (**Fig. 8A-C**) revealed significant synergistic interactions between dasatinib and capivasertib in all three cells, with mean synergy score of 12.42 in MCF10-PTEN-KO1 cells, 11.73 in HCC1937 and 10.99 in SPAC-1-L cells, each with highly significant p-values, considering that a synergy score larger than 10 suggests the interaction between two drugs is likely to be synergistic.(**Fig. 8A-C**). We further performed western blot analysis to examine the phosphorylation levels of Src pY416 and GSK3 α/β pS21/9 to confirm the inhibition of Src and AKT signaling. Additionally, the apoptotic markers, cleaved PARP, and cleaved Caspase-3 proteins were observed to be more prominent in cells treated with the combination compared to each drug alone, suggesting that dual inhibition of Src and AKT may enhance apoptotic effects (**Fig. 8D-F**). This highlights the therapeutic potential of combining dasatinib and capivasertib to achieve superior efficacy compared to individual treatments in PTEN-deficient cells.

**Figure 8.**
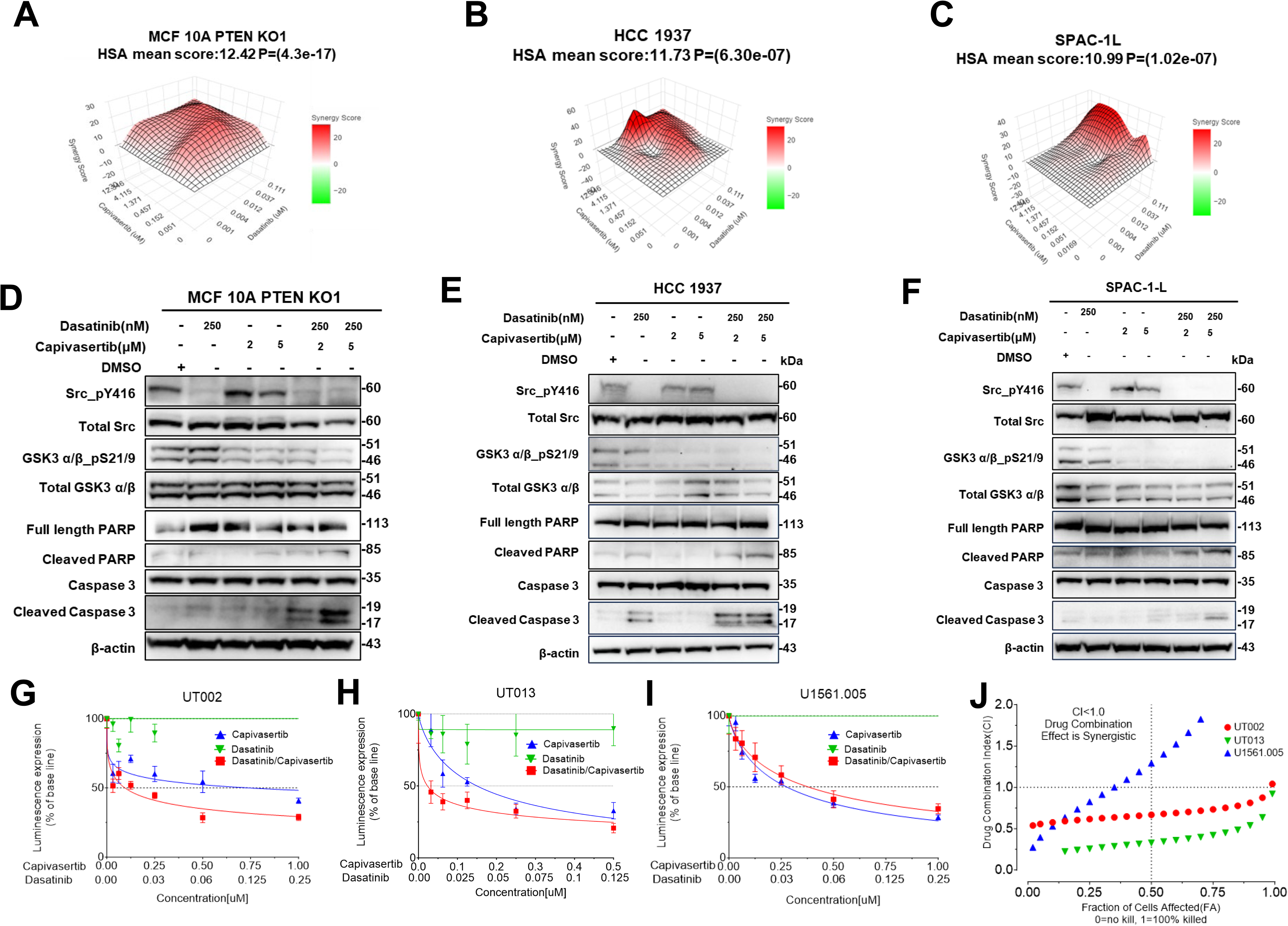
AKT inhibitor and Src inhibitor synergistically suppress cancer cells with PTEN loss. (**A-C**) Synergy Analysis of Dasatinib and Capivasertib in PTEN-Deficient Cells. 3D synergy plots for Dasatinib and Capivasertib combinations in PTEN-deficient cell lines (A) MCF10A PTEN KO1, (B) HCC1937, and (C) SPAC-1L. High synergy scores (red regions) indicate strong synergistic effects in all cell lines. (**D-F**) Western blots showing Src and AKT pathway inhibition (Src_pY416 and GSK3 α/β_pS21/9) and increased apoptotic markers (cleaved PARP and Caspase-3) with Dasatinib and Capivasertib combination in (D) MCF10A PTEN KO, (E) HCC1937, and (F) SPAC-1L cells. (**G-I**) Dose-Response of Combination Treatment in PTEN-Deficient PDX Models. Dose-response curves for the treatment of Dasatinib, Capivasertib or combination in PTEN-deficient endometrial cancer PDX models (**G**) UT002.HS3.A.4.L.L, (**H**) UT013.HS2.0.0.B, and (**I**) U1561.005.HS3.J.4.G, showing enhanced suppression of cell viability with combination treatment in *ex vivo* 3-D culture. (**J**) Combination Index (CI) plot illustrating the synergistic effects of the drug combination in PTEN-deficient endometrial cancer PDX models (UT002, UT013, and U1561.005). The x-axis represents the fraction of cells affected (FA), The y-axis shows the CI values, with CI < 1 indicating synergy. Drug combinations exhibit synergistic effects (CI < 1) in UT002 (red circles) and UT013 (green triangles) models, while U1561.005 (blue triangles) shows no significant synergy at most FA levels.

To further validate the synergistic effect of the combination treatment, we utilized three endometrial cancer patient-derived xenograft (PDX) tumor models with PTEN deficiency (**Supplemental Data 9)**. Two of the models, UT002 and UT013, were established from primary surgery samples of patients with high-grade endometrial carcinosarcoma. UT002 harbored a PTEN loss-of-function missense mutation (Y155C) located within the phosphatase domain critical for PTEN stability and lipid phosphatase activity (55, 56). UT013 carried R130G and N276 frameshift mutations. Arginine 130, situated near the core catalytic domain, is a known mutation hotspot, accounting for approximately 20% of PTEN mutations in endometrial cancer (57). The third PDX model, U1561.005, was derived from a patient with high-grade endometrioid cancer and also carried the R130G mutation. In 3D *ex vivo* cultures, all three PDX models were treated with dasatinib, capivasertib, or their combination and assayed for the cell viability. The combination therapy demonstrated significantly lower cell viability in UT002 and UT013 models compared to either drug alone (**Fig. 8G-I**). No synergy was observed in U1561.005 PDX model. Interestingly, dasatinib alone exhibited minimal growth suppression in all three PDX models. Synergy analysis confirmed that dual inhibition with dasatinib and capivasertib synergistically reduced cell viability in UT002 and UT013 PDX models (**Fig. 8J**). These findings underscore the potential of combining dasatinib and capivasertib as a promising therapeutic approach to effectively target PTEN-deficient subsets of endometrial cancer.

## Discussion

PTEN is a potent tumor suppressor (3, 58), serving as a critical player in the oncogenic PI3K signaling pathway (4). Additionally, it has PI3K-independent functions associated with its protein phosphatase- and nuclear activities (59), underscoring its multifaceted role in cellular processes. Yip and colleagues performed TiO_2_-enriched phosphoproteomics in mouse embryonic fibroblasts (MEFs) with *Pten* mutations to investigate the PI3K-independent functions of PTEN as lipid- or protein phosphatase (60). In another study, the TiO_2_-enriched phosphoproteome of MEFs with oncogenic *Pik3ca* mutation and *Pten* deletion were compared(61). However, there remains a gap in unbiased, systematic proteome and phosphoproteome profiling in human epithelial cells, where PTEN loss is critically involved in cancer development. In this study, we set out to explore *PTEN* knockout-induced changes in the total proteome, IMAC-enriched phosphoproteome (mainly phosphoserine/threonine), and pTyr-1000 enriched phosphotyrosine proteome in a nontumorigenic human mammary gland epithelial cell line, MCF10A.

Pathway enrichment analysis of our proteomics data revealed that PTEN loss influences cellular signaling events beyond the canonical PI3K pathway. The biological disruptions observed in our study align with previously reported oncogenic pathways altered by PIK3CA mutations, including cytoskeleton rearrangement, cell cycle regulation, MAPK, AKT/mTOR, and ERBB2 signaling (24). Gene set enrichment analysis (GSEA) further reinforces the established connection between PTEN and p53, as well as cyclin D1 (13, 15), highlighting PTEN’s potential role in regulating a diverse array of transcription factors (62, 63). Additionally, our pathway analysis suggests that PTEN loss enhances central carbon metabolism, including cytosolic glycolysis, the TCA cycle, and oxidative phosphorylation in mitochondria.

More importantly, our phosphoproteomics analysis uncovered that beyond the canonical serine-threonine kinase signaling pathways regulated by the PTEN-PI3K-AKT signaling cascade, PTEN loss can also induce global activation of tyrosine kinase signaling networks (**Fig. 5E-F**). Accordingly, multiple RTKs, including five RTKs (EGFR, EphA2, EphB2, EphB4, and MET) and six non-RTKs (FRK, YES1, PTK2, PTK2B, JAK2, and TYK2), were found be hyperphosphorylated in cells with PTEN loss (**Fig. 5L, 6A**). PTEN shares sequence homology with protein tyrosine phosphatases that directly dephosphorylate tyrosine sites (58). PTK2 (FAK), PTK6, Src, and FYN were previously reported to be dephosphorylated and suppressed by PTEN(7, 59, 64, 65). Although only PTK2 hyperphosphorylation was directly observed in our phosphoproteomic analysis, our kinase enrichment analysis suggested that Src and FYN were among the activated upstream tyrosine kinases in addition to PTK2 that contributed to the global elevation of tyrosine phosphorylation in cells with PTEN loss (**Fig. 5I-K**).

Notably, several Ephrin receptor tyrosine kinases, EphA2, EphB2 and EphB4, were hyperphosphorylated in PTEN-KO cells. In addition, EphA2 mRNA and protein were also upregulated (**Fig. 3E and J**). EphA2 overexpression, activation and its association with poor patient survival outcomes have been frequently reported in a wide range of cancer types (44). Exploring the TCGA dataset from cBioPortal (38), we found that breast cancer patients with PTEN low and EphA2 high expression have the worst prognosis among the patients with different levels of PTEN and EphA2 expressions (**Fig. 6B**). Using different cancer cell lines and genetic manipulation methods, we confirmed that PTEN loss leads to the EphA2 upregulation across different cancer types (**Fig. 6C-E**). Interestingly, inhibiting AKT with AKT-specific inhibitors, MK2206, ipatasertib or capivasertib, could not reduce EphA2 expression (**Fig. 6M-N**). Instead, suppression Src with Src siRNA or the Src inhibitor dasatinib reduced EphA2 expression in cells with PTEN loss (**Fig. 7D-I**). These data indicated that activation of Src kinase, rather than canonical AKT kinase signaling, is critical in upregulating EphA2 in cancer cells with PTEN loss. A corollary of these results is that targeting AKT alone in PTEN-deficient or PTEN-mutant cancers might be insufficient due to the activation of noncanonical tyrosine kinase signaling pathways. Furthermore, PTEN loss has been reported to contribute to AKT inhibitor resistance in multiple cancers(66). In the LOTUS trial, which evaluated the efficacy of the AKT inhibitor ipatasertib combined with paclitaxel in patients with metastatic triple-negative breast cancer, patients with PTEN-low tumors did not experience significant benefits, indicating potential resistance to AKT inhibition in the context of PTEN loss (66). Collectively, these observations suggest that AKT as well as EphA2 or Src might need to be inhibited to have maximal impact on PTEN null cells. To investigate the potential synergy of combinate treatment with AKT and Src inhibitors, we treated PTEN-deficient breast and endometrial cancer cell lines with dasatinib and capivasertib. Our results demonstrated that targeting both AKT and Src kinases can significantly reduce cancer cell proliferation and substantially induce apoptosis (**Fig. 8A-F**). Furthermore, in endometrial cancer PDX models, combination treatment with dasatinib and capivasertib showed enhanced effectiveness compared to single-agent treatments (**Fig. 8G-H**), suggesting that dual-targeted therapies could offer a promising approach to overcome PTEN-loss-driven resistance and improve therapeutic efficacy in cancer treatment.

In summary, our quantitative proteomics and phosphoproteomics analyses revealed that PTEN loss activates diverse signaling pathways involved in tumor initiation and progression. In addition to the canonical serine/threonine kinase-based PI3K-AKT pathway, PTEN loss also activates a wide range of tyrosine kinase-mediated signaling networks. Our study provided novel evidence that PTEN loss upregulates EphA2 receptor tyrosine kinase (RTK) expression through the activation of Src tyrosine kinase. Furthermore, we demonstrated that dual inhibition of AKT and Src kinases, both activated as a consequence of PTEN loss, significantly suppressed tumor cell growth and induced apoptosis. These findings underscore the therapeutic potential of combining AKT and Src inhibitors in PTEN-deficient cancers, addressing the limitations of AKT inhibition alone and offering a promising strategy to overcome resistance associated with PTEN loss.

## Abbreviations

PTEN: Phosphatase and tensin homolog
PI3K: Phosphatidylinositol-3-kinase
PIP2: phosphatidylinositol 4,5-bisphosphate
PIP3: phosphatidylinositol 3,4,5-bisphosphate
MS: mass spectrometry
RTK: Receptor tyrosine kinase
KO: Knockout
KD: Knockdown
PDX: Patient-derived xenograft
EGF: Epidermal growth factor
EGFR: Epidermal growth factor receptor
MEF: mouse embryonic fibroblast
IMAC: Immobilized metal affinity chromatography
TCA: tricarboxylic acid cycle
GSEA: Geen set enrichment analysis
TMT: Tandem mass tag
PCA: Principal Component Analysis

## Acknowledgements

This work was supported by the by the Mayo Clinic Breast Cancer Specialized Program of Research Excellence (SPORE) (P50CA116201) Developmental Research Program Award, Mayo Clinic Ovarian Cancer Specialized Program of Research Excellence (SPORE) (P50 CA136393) Developmental Research Program Award, a DOD Breast Cancer Research Breakthrough Award (BC181309) and Minnesota Ovarian Cancer Alliance Research Award to X.W. This work was supported by grants from NCI’s Clinical Proteomics Tumor Analysis Consortium (U01CA271410) and Mayo Clinic Comprehensive Cancer Center grant (P30CA15083) to A. P.. B.H.P. acknowledges support from Susan G. Komen, the Breast Cancer Research Foundation and NIH/NCI grants P30CA06485, P50CA098131 and R01CA289528.

We thank Dr. Scott Kaufmann for providing ovarian and endometrial cancer cell lines, providing helpful advice, and commenting on the manuscript.

## Data availability

The MS data were deposited to the ProteomeXchange Consortium via the PRIDE partner repository and are available with the accession number PXD057520. Reviewers can access the dataset by logging in to the PRIDE website using the following account details:

Username

Password: juPz5t92uxVs

## Declarations

### Ethics approval and consent to participate

All procedures were conducted in accordance with Animal Welfare Regulations and were approved by the Institutional Animal Care and Use Committee (IACUC) at the Mayo Clinic.

### Consent for publication

All authors have reviewed and approve the final manuscript and consent to its publication.

### Authors’ contributions

X.W., A.P., H. Z., Q.W., X.K., H. S. and L. F. conceived ideas, designed experiments, interpreted results, and wrote the manuscript. X.W. Q.W., X.K. H. S., L. W., L. L, X. H. R. C. M. K. H. K. and J. W., performed cell biology and molecular biology studies. X.H. and J. S. W. performed PDX studies. X.W., Q. W., L. W. K. M. and S. R. performed MS-based proteomic analysis. T. L. L and B. H. P. preformed PTEN KO and provided the PTEN-KO cells.

