## Supplementary Figures for "Proteomic Analysis of PTEN-Deficient Cells Reveals Src-Mediated Upregulation of EphA2 and Therapeutic Potential of Dual Inhibition"

**Affiliations:**

**Fig. S1**

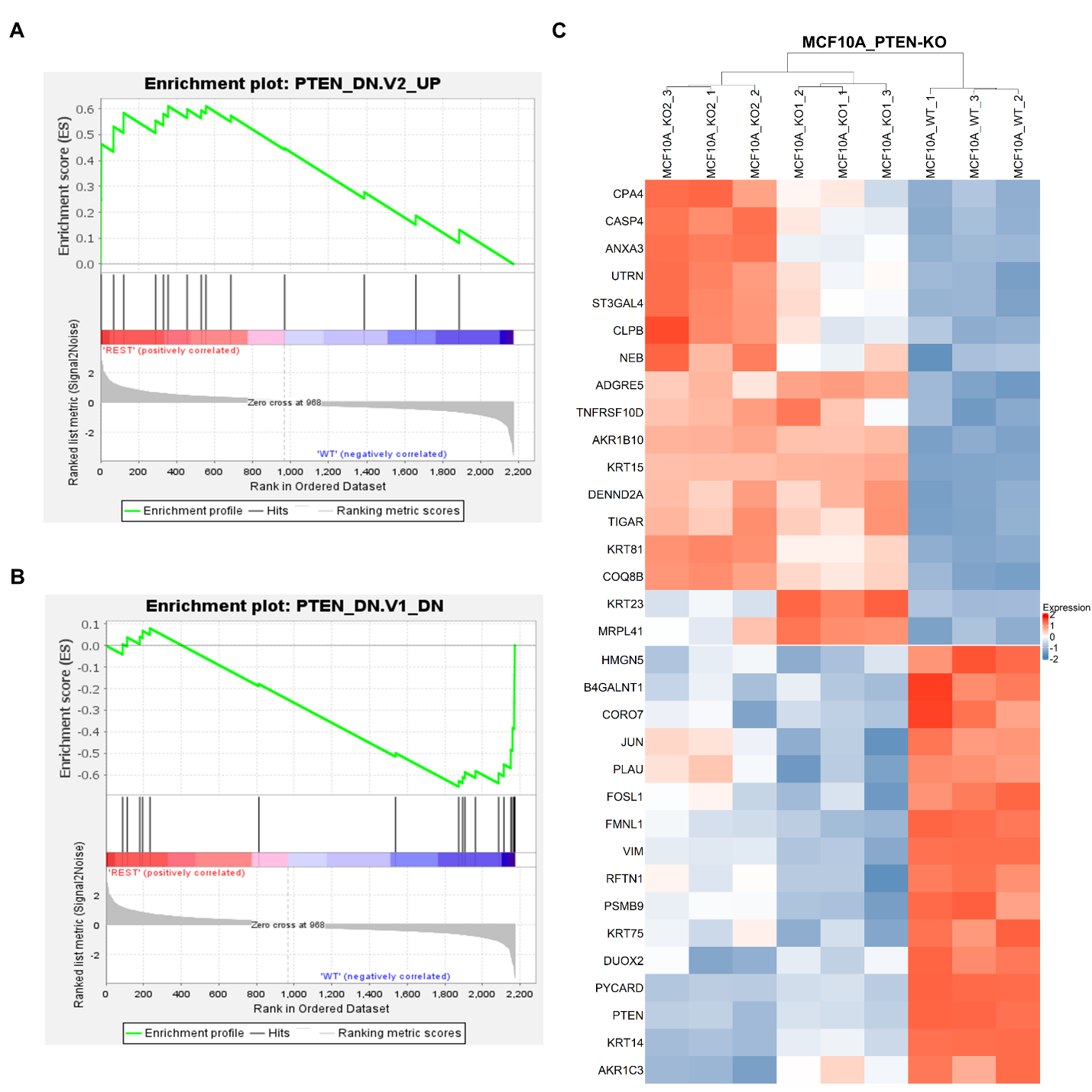

***Fig. S1.*** **Enrichment plots of the enriched PTEN gene sets and their gene set members in GSEA.** The gene set members in the analyzed and enriched gene sets “PTEN_DN.V2_UP” **(A)** and “PTEN_DN.V1_DN” **(B)** that were included in the proteomic dataset were plotted in the heatmap **(C)**. Red: protein upregulation; Blue: protein downregulation.

**Fig. S2**

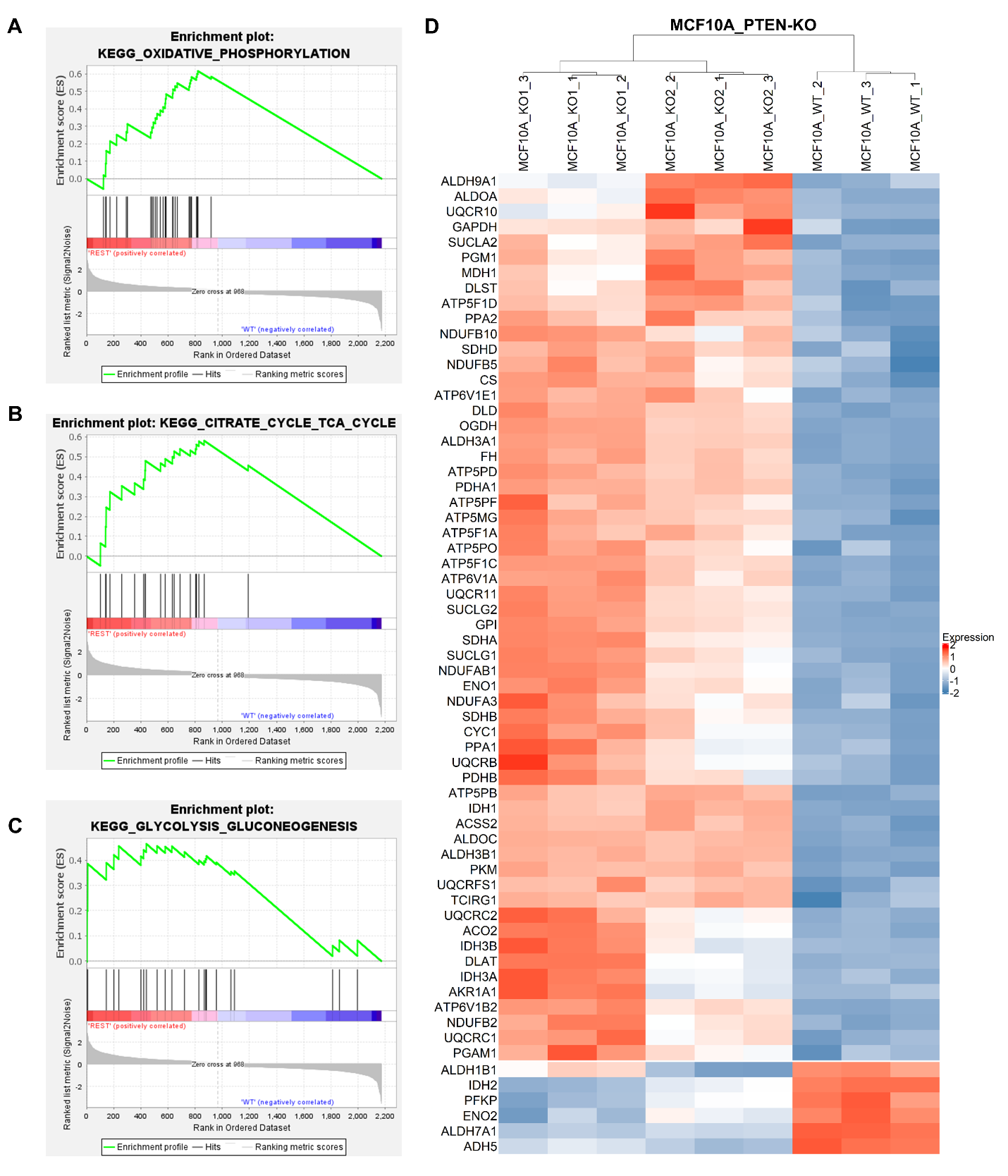

***Fig. S2.*** **Enrichment plots of the enriched central carbon metabolic gene sets and their gene set members in GSEA.** The gene set members in the analyzed and enriched central carbon metabolism gene sets “KEGG_OXIDATIVE_PHOSPHORYLATION” **(A)**, “CITRATE_CYCLE_TCA_CYCLE” **(B)**, and “KEGG_CLYCOLYSIS_GLUCONEOGENESIS” **(C)** that were also included in the proteomic dataset were plotted in the heatmap **(D)**. Red: protein upregulation; Blue: protein downregulation.

**Fig. S3**

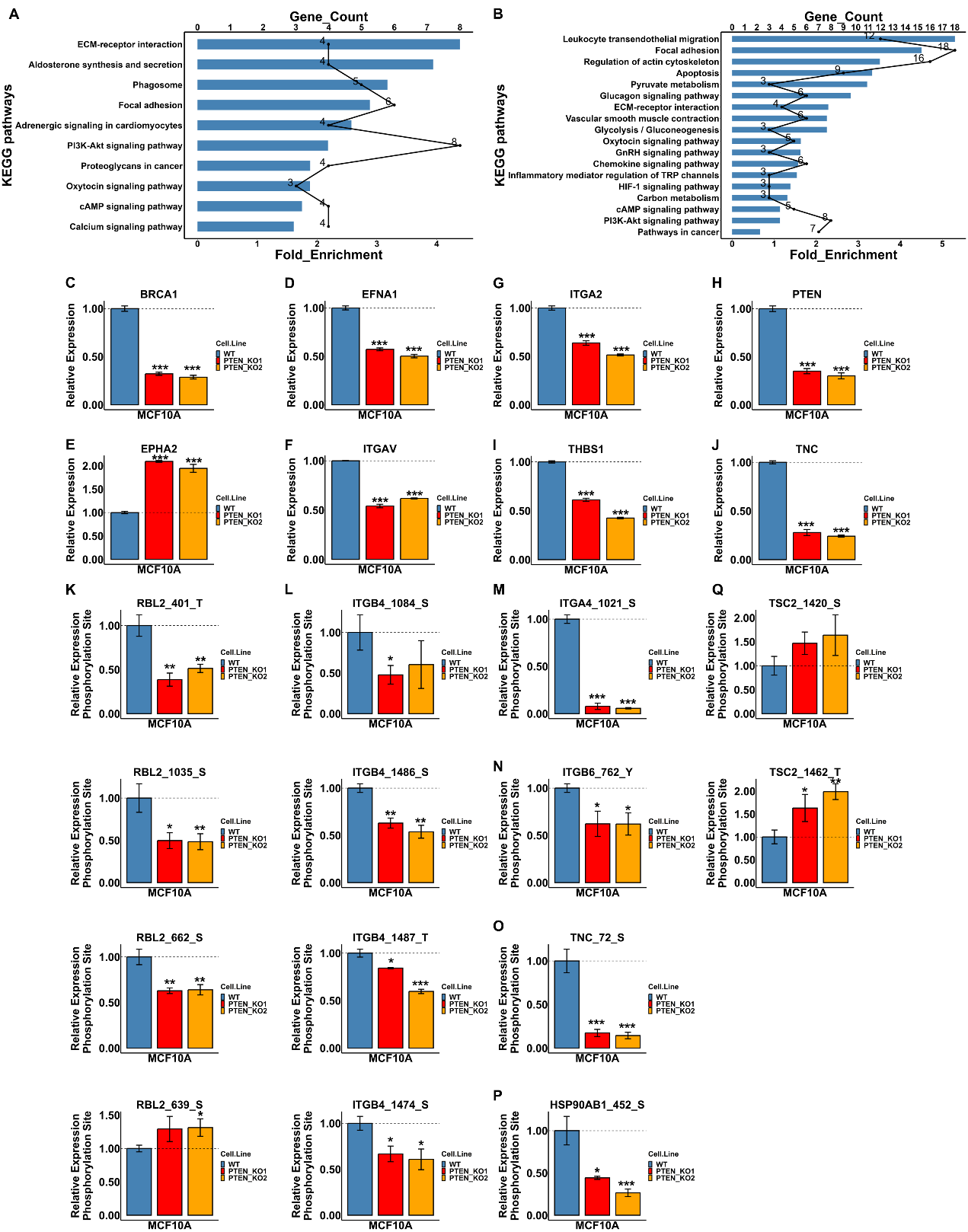

***Fig. S3.*** **KEGG pathway enrichment and the identified significantly dysregulated PI3K-AKT signaling components**. **(A-B)** Significantly changed proteins **(A)** and phosphoproteins **(B)** (Student’s T-test p value < 0.05; fold change >= 1.5) were analyzed for KEGG pathway enrichment in DAVID (Version: Dec 2021). Default parameter setup on DAVID was applied. Bar: fold enrichment; Line: gene count in each enriched pathway. **(C-Q)** Gene components in the PI3K-AKT KEGG signaling pathway were selectively plotted for their protein and/or phosphorylation levels detected in this study. Protein/phosphoprotein expression levels of *PTEN* knockout were normalized to the wide type. Two sample Student’s T-test were performed for each knockout clone compared with the wide type. *: p value < 0.05; **: p value < 0.01; ***: p value < 0.001.

**Fig. S4**

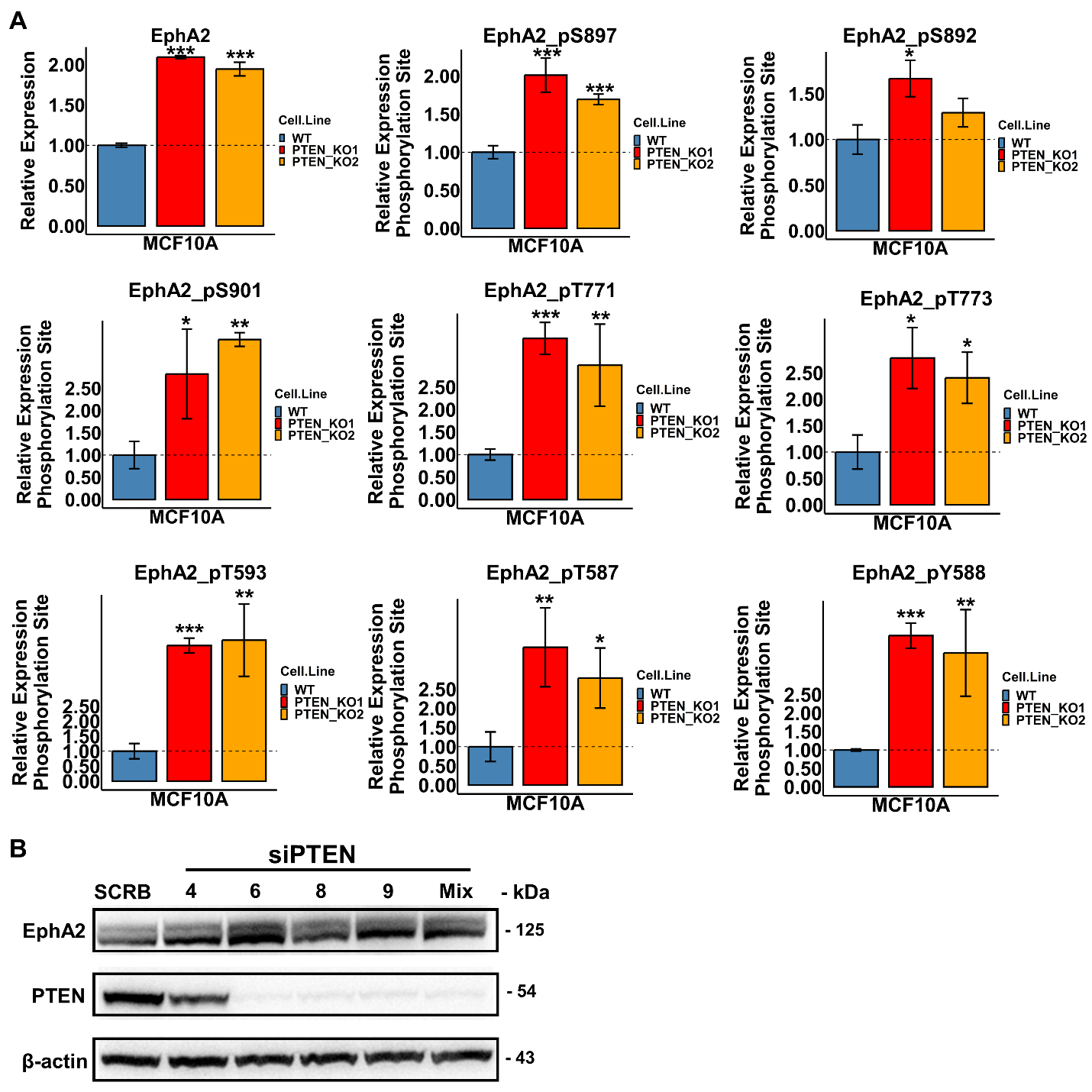

***Fig. S4.* Functional assays for PTEN and EphA2. (A)** Total protein expression of EphA2 and levels of the phosphorylated sites detected on EphA2 by IMAC- and p-Tyr-1000 enrichment were plotted. **(B)** Knockdown efficiency of the siRNAs targeting PTEN.

**Fig. S5**

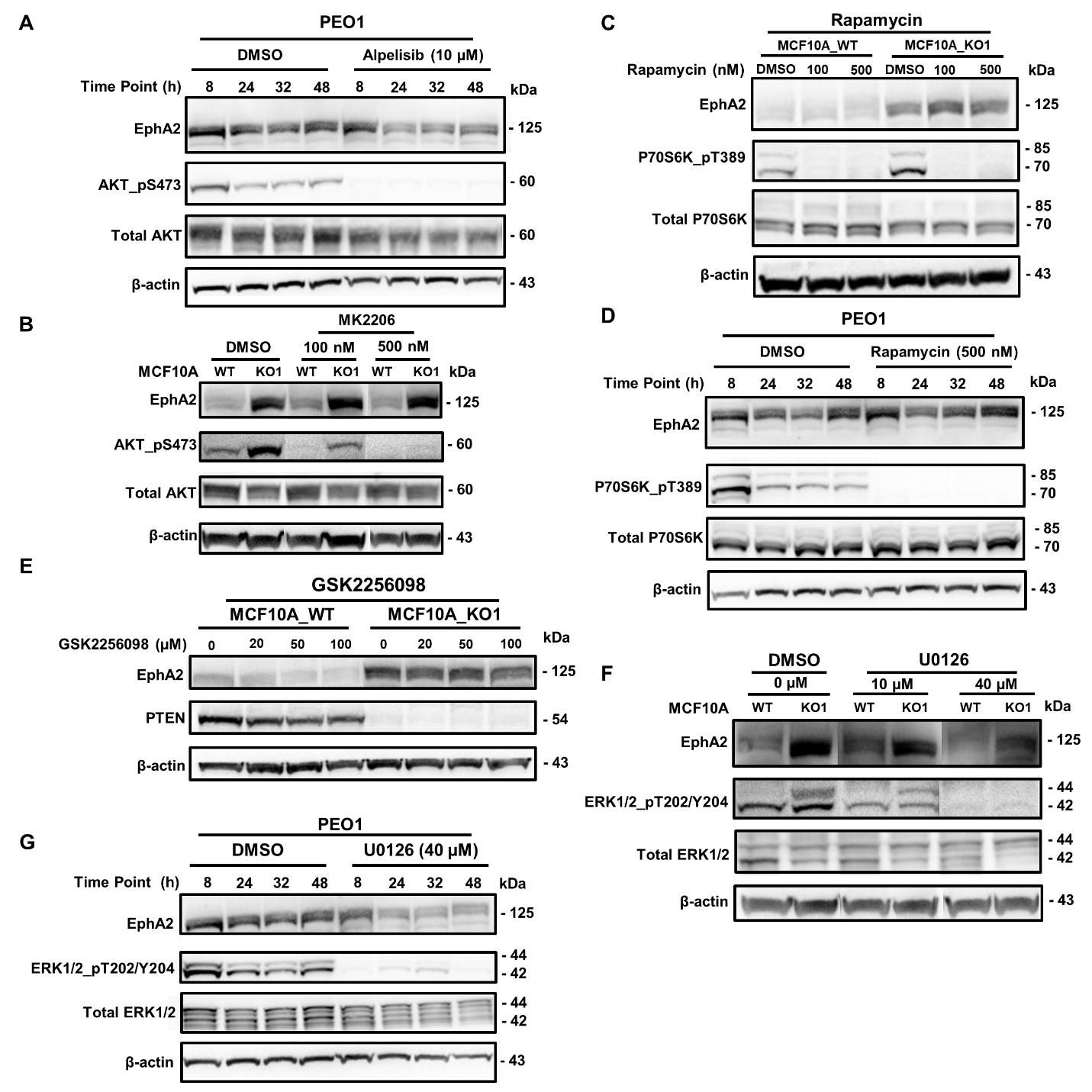

***Fig. S5.* The mechanism of PTEN in regulating EphA2 protein expression. (A)** Time series of the PI3K inhibitor Alpelisib (10 µM) treatments in PEO1. **(B)** MCF10A wide type and PTEN knockout cell clone 1 were treated with the AKT inhibitor MK2206 (100 nM and 500 nM). **(C)** MCF10A wide type and PTEN knockout cell clone 1 were treated with the mTOR inhibitor Rapamycin (100 nM and 500 nM). **(D)** Time series of the mTOR inhibitor Rapamycin (500 nM) treatments in PEO1. **(E)** MCF10A wide type and PTEN knockout cell clone 1 were treated with the FAK1 inhibitor GSK2256098 (20 µM, 50 µM and 100 µM). **(F)** MCF10A wide type and PTEN knockout cell clone 1 were treated with the MEK inhibitor U0126 (10 µM and 40 µM). **(G)** Time series of the MEK inhibitor U0126 (10 µM and 40 µM) treatments in PEO1.
